# LongevityBench: Are SotA LLMs ready for aging research?

**DOI:** 10.64898/2026.01.12.698650

**Authors:** Alex Zhavoronkov, Denis Sidorenko, Vladimir Naumov, Stefan Pushkov, Diana Zagirova, Vladimir Aladinskiy, Derya Unutmaz, Alex Aliper, Fedor Galkin

## Abstract

Aging is a core biological process observed in most species and tissues, which is studied with a vast array of technologies. We argue that the abilities of AI systems to emulate aging and to accurately interpret biodata in its context are the key criteria to judge an LLM’s utility in biomedical research. Here, we present LongevityBench — a collection of tasks designed to assess whether foundation models grasp the fundamental principles of aging biology and can use low-level biodata to arrive at phenotype-level conclusions. The benchmark covers a variety of prediction targets including human time-to-death, mutations’ effect on lifespan, and age-dependent omics patterns. It spans all common biodata types used in longevity research: transcriptomes, DNA methylation profiles, proteomes, genomes, clinical blood tests and biometrics, as well as natural language annotations. After ranking state-of-the-art foundation models using LongevityBench, we highlight their weaknesses and outline procedures to maximize their utility in aging research and life sciences.

## 1. Introduction

Aging is among the most information-rich and universal phenomena in biology. It unfolds across all tissues, involves every organizational layer, and can be observed in a variety of species. Building a digital model of aging is thus among the most challenging tasks in modern biology, since it would need to operate in a wide variety of contexts. A model that genuinely understands aging should recognize how molecular regulation shifts with age, connect low-level measurements to organism-level phenotypes, and be able to interpret the footprints of aging in omics and other biodata types. These requirements make aging biology an ideal domain for evaluating whether AI systems operate from coherent internal representations of life or rely on superficial correlations and fact memorization.

Deep learning has been applied to aging research for over a decade. Early works in this field focused on measuring its intensity within the aging clock paradigm and aging biomarker discovery [1]. Throughout the 2010s, neural networks revealed aging patterns in DNA methylation (DNAm) [2], gut flora composition [3], mRNA expression [4], facial photography [5], clinical blood tests [6], and psychological assessments [7]. These models found practical applications in batch effect correction [8], target discovery [9], and clinical trial design [9]. Subsequent architectures moved beyond description toward data generation by learning joint representations across data types to identify compounds that induce desired pro-longevity molecular changes [10, 11]. Most recently, the transformer-based systems have enabled multi-modal omics comprehension and simulation of aging processes across species [12–14].

The trajectory of dedicated biodata-focused AI development is now intertwined with the progress in the general-purpose large language models (LLMs). Commercial LLMs, such as ChatGPT, Claude, Grok, Gemini are now deeply embedded in research workflows, increasing scientists’ productivity as academic authors [15]. AI applications help scientists with routine tasks, such as writing text and code or finding references, and ultimately save their time for more creative activities. Some envision an even deeper integration via agentic systems that can handle end-to-end research projects [16]. However, skepticism persists and concerns about the negative effects of overreliance on AI are often expressed on professional forums and in private discussions, though peer-reviewed critiques rarely propose concrete remedies [17–19].

We argue that the research community’s worries can be attenuated by a disciplined approach to model evaluation. Benchmarks provide an empirical foundation for calibrating user trust: a model that fails to correctly interpret standard biodata cannot be trusted as a research companion, regardless of how fluently it communicates. The field of aging biology offers a wide range of tests that can be used to judge an LLM’s grasp of the biological ground truth and ability to reason, and thus, evaluate its fitness for taking part in the scientific process. Despite the importance of aging research and the growing adoption of LLMs in biomedical workflows, no systematic benchmark exists to evaluate whether these models can reliably interpret biodata in the context of aging. A benchmark set covering varied data types, biological scales, and question formats could be used as a test for the presence of a coherent internal model of life, making it not just a way expose LLMs competence gaps, but also a development target. By steering model training toward mastery in all challenges presented by aging research, we can achieve models operating based on genuine understanding of the biological reality rather than superficial pattern-matching.

To this end, we present LongevityBench, a comprehensive evaluation framework designed to assess whether foundation models can operate competently in the aging biology domain. The benchmark encompasses tasks requiring models to predict survival from clinical records, infer age from omics profiles, determine how genetic perturbations affect lifespan, and identify age-related expression patterns. It spans the major data modalities of longevity research: transcriptomics, epigenetics, proteomics, clinical biochemistry, and genetic intervention studies. Importantly, each task is presented in multiple question formats, allowing us to distinguish whether a model can operate effectively independent from query semantics.

LongevityBench serves two purposes. First, it provides a leaderboard for state-of-the-art (SotA) LLMs, helping researchers select appropriate tools for their aging-related analyses. Second, it establishes evaluation infrastructure for our ongoing development of research-grade biological AI. The benchmark constitutes a module within our Multi-Modal AI (MMAI) Gym for Science (Figure 1), a pipeline designed to enhance foundation models’ utility in life science research. While this paper focuses on evaluating existing commercial models, the approaches we have demonstrated in LongevityBench provide a foundation for future works aimed at training more advanced AI systems rather than measuring their performance.

**Figure 1.**
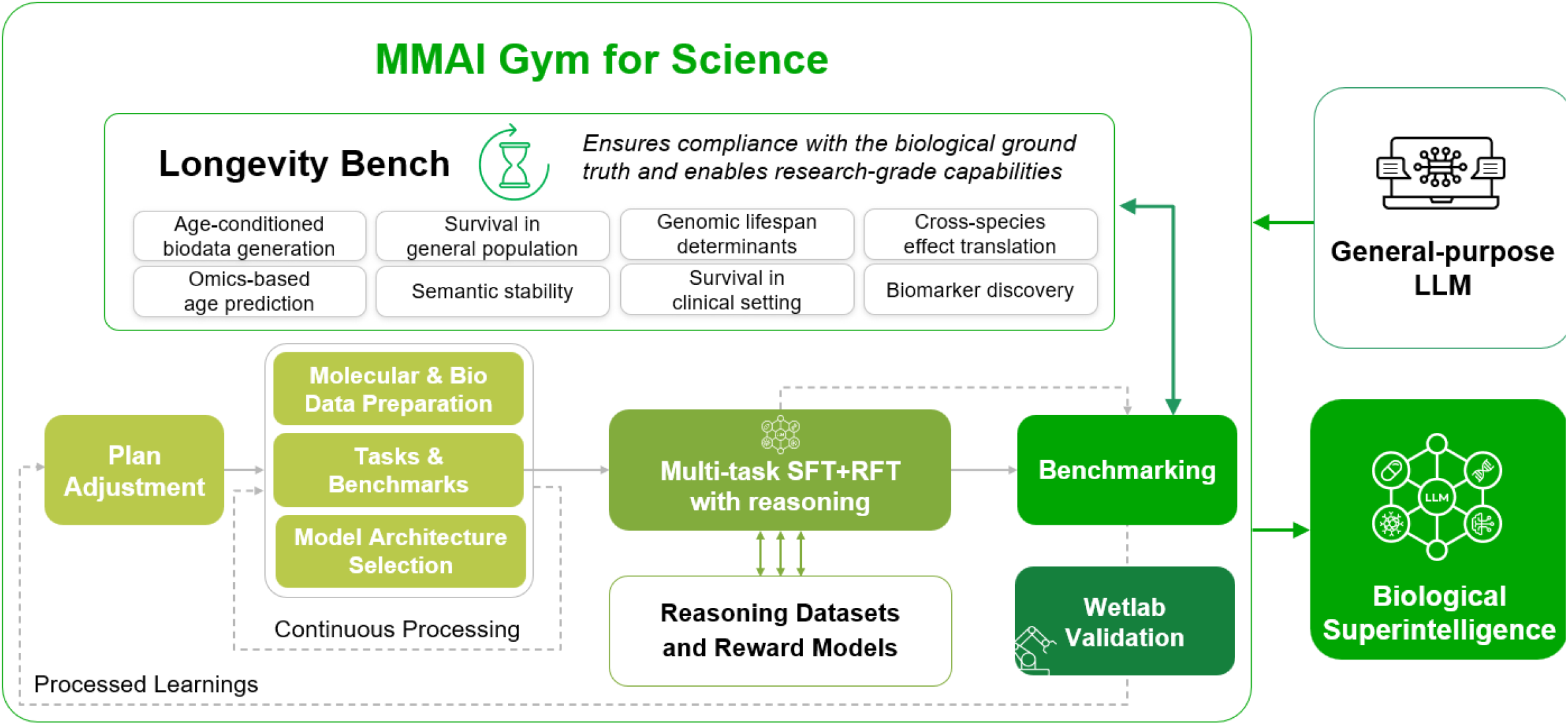
High-level scheme of an LLM training in the Multi-Modal AI (MMAI) Gym for Science. The gym is an iterative pipeline which leverages a variety of deep learning techniques, such as supervised fine-tuning (SFT) and reinforcement fine-tuning (RFT) based on carefully annotated biological and chemical datasets to enhance LLMs’ alignment with the empirical ground truth. In this pipeline, LongevityBench serves a crucial role by grounding the training process with a set of tests aimed at evaluating a model’s understanding of aging biology.

## 2. Materials and Methods

### 2.1. Benchmark Datasets

To create LongevityBench, we used multiple publicly available data sources to compose structured prompts in formats that may be processed by LLMs and enable straightforward metrics calculations. The datasets were selected to contain relevant aging-related annotations, which include chronological age, group lifespan, and survival information. Each data source was prepared in several alternative ways that differ by either the depth of the context provided to the model in the prompt or the format in which the model is instructed to provide the answer. The presented collection features prompts that request the model to carry out the regression task (e.g. specify the exact time-to-death), binary classification (e.g. will an individual survive >10 years), pairwise selection (e.g. which individual will live longer) with binary or ternary options, depending on whether tied pairs are included in the collection, and multiple choice (e.g. will the individual survive <5, 5-10, >10 years). For all multiple-choice formats, the order of presented options was randomized. All prompts were formatted as XML-like structures with tags separating different context sections, enabling easy control over the amount of context an LLM received during testing. All prompts were supplemented with a “`<follow_up>`” section to enable manual curation and sanity checks, which was removed during benchmarking. See **“Supplementary materials”** for prompt samples.

See below the details of data collection, partition, and processing for each category with sample prompts for each supported format. All data sources were used only partially, to enable later independent tests and experiments with the unused entries.

#### General population survival

We used the two oldest waves (1999, 2001) from the NHANES-IV dataset to compose the prompts in this collection.

For the binary format, we asked LLMs if an individual survived for at least 10 years post-examination. The benchmark contained 8976 prompts with individuals balanced for sex and age group, among them 1356 confirmed deceased with 194±61 survival time. The natural text annotations for the prompts were prepared using 54 variables from the harmonized NHANES dataset (CC BY 4.0 licence) [20] describing demographic parameters of each individual, details of their health history and physical fitness. The blood biomarker data to supplement the prompts was obtained through the `pynhanes` package (MIT licence) [21]. The mean number of blood markers per sample was 28, in the training set and 29 in the test set.

Apart from the binary formats, we prepared pairwise time-to-death (TTD) comparisons with 15,542 pairs of NHANES participants with 4697 unique individuals. Finally, wo types of prompts were created for singular TTD prediction: (i) binned with options specified as “0-5 years”, “5-10 years”, “10-15 years”, and “15+years”, or (ii) regression format, in which an LLM was required to predict the exact TTD of a participant in months.

#### Aging trajectories

We used the Open Genes [22] database (MPL-2.0 licence) to compose prompts that asked an LLM to assess the directionality of a gene’s expression change in human tissues. Based on this dataset, we prepared 647 prompts, featuring diverse tissues, such as muscle, CD8+ T-cells, frontal cortex and others. The prompts represent mostly mRNA-level expression changes and have a balanced proportion of genes with an upward or downward aging trajectory (54% vs 46%). Among the 1953 unique genes found in the Open Genes database, 157 were annotated with findings in model organisms, such as mouse, rats, and others. The prompts contain additional sections providing context about a masked gene’s function, based on Gene Ontology (GO), and human tissue localization, based on Protein Atlas.

#### Multi-mutant lifespan

We used the Synergy Age [23] (CC BY 4.0) database to prepare prompts featuring genetic interventions into model organisms with known effect on the lifespan. 48 *Drosophila melanogaster* and 16 *Mus musculus* experiments from the database were formatted as text prompts. The pairwise prompts required an LLM to assess whether a single-mutant or a double-mutant organism had a longer lifespan. For 41% of prompts, the double mutant was the correct answer. For the regression prompt format, the model was asked to provide the exact percentage increase in lifespan for a double-mutant organism compared to the wild type (WT). In this collection, 39% of double mutants had increased life duration. As part of our ablation study, the pairwise prompts were prepared in the minimal format containing only lifespan durations of single mutants as extra context, or in the extended format featuring additional GO annotations and experimental context pulled from source publications.

#### Cancer survival

We used TCGA Open Access data, as defined in the NIH GDS and NCI GDS policies, to prepare prompts featuring RNAseq findings, patient diagnosis and treatment information, to predict patients’ progression-free survival interval (PFS). The KICH, LUAD, and SKCM cohorts were used to derive text prompts.

All prompts were formatted as pairwise choice with randomly selected two patients with the same tumor type. Each prompt is supplemented with processed RNAseq data. For each patient we ran single-sample gene set enrichment analysis (ssGSEA) against the Reactome pathway database to select the most characteristic pathways for each patient, based on normalized enrichment scores (NES) statistics. During the ssGSEA stage, we used fold changes between a patient’s normal and malignant tissues to create preranked lists. The prompts contained up to 5 pathways with the most dissimilar NES between patients in a pair, and up to 5 pathways with extreme NES scores across all patients (<5^th^ or >95^th^ percentiles of the total distribution). Additionally, the prompts displayed up to 10 most highly expressed genes for the featured pathways with a positive NES, or 10 genes with the lowest expression values from the pathways with a negative NES. By avoiding exact expression values and focusing on pathway-level annotations, we thus managed to present informative and compact RNAseq context for tumor environments. In the end, the prompt collection contained 424 Reactome pathways and 3,740 unique genes.

#### Methylation aging

We used a collection of Illumina Human Methylation studies released on Gene Expression Omnibus (GEO) described in [2] to create prompts aimed at measuring the ability of LLMs to process epigenetic signals with minimal processing into chronological age predictions. The DNAm benchmark prompts contained samples from the following GEO studies: GSE102177, GSE103911, GSE105123, GSE107459, GSE107737, GSE112696, GSE20067, GSE34639, GSE37008, GSE59065, GSE61496, GSE79329, GSE87582, GSE87640, GSE98876, GSE99624. When available, prompts featured information about samples’ sex, associated diseases, smoking, BMI, pregnancy, and alcohol use. The prompts were prepared in three formats. For the multiple-choice format, LLMs were asked to choose the by-decade age group of a single sample, ranging from “0-9” years to “90-99”, and “101+”. The resulting collection contains 500 prompts. The same samples were also used to prepare the regression-formatted prompts, in which LLMs were instructed to predict the exact age in years. Finally, we prepared an equal number of pairwise prompts, for which LLMs were instructed to select which of the two samples was older. The prompts for this format contained 668 unique subjects. Cross-study comparisons were enabled for the pairwise prompts.

To represent the DNAm information in the prompts, we used beta-values from 1,000 CpG sites used by the DeepMAge aging clock [2]. For the single-profile formats, we selected 75 most and 75 least methylated sites, and for the pairwise prompts we showed 150 most dissimilar sites between two profiles. To reduce dependence on Illumina’s internal site reference numbers, we referred to them by genomic locations and probe target sequences instead, as described in Infinium Human Methylation platforms’ manifest files. All coordinates were stated in the hg38 human genome assembly convention. For sites located near or within known genes, brief annotations of these genes were included, as per the platforms’ manifest files.

#### Transcriptomic aging

We used open access data from GTEX, governed by the <u>GTEx Portal Data License</u>, to instruct the models to predict the age of multi-tissue human samples. We used 162 subjects to compose the prompts. The data scope in GTEX contained 31 different tissues six age groups binned by decade (from “20-29” to “70-79”).

The prompt collection includes multiple choice binned age prediction, pairwise binary and generative formats. In total, these formats contained 1,200 prompts. For the pairwise task, the binary format implied pairs of samples in which one sample was always older than the other. No cross-tissue pairs were included in the pairwise prompt collections.

In the pairwise setting, top-100 expressed genes were presented for each sample and supplemented with contrastive ssGSEA finding, similar to the approach used in the cancer survival task. In the binned age prediction, 200 genes with numeric intensity values from each decile were shown, alongside the number of gene with nTPM<0.5 (marked as “not detected”) and top-200 highly expressed genes. For the generative task, we presented LLMs with random 50 genes from among top-100 expressed genes in a sample and instructed the models to generate the remaining 50 in any order.

#### Proteomic aging

To assess if LLMs can pick up the aging signal in proteomic data, we used the <u>PAD000022</u> Olink Explore 3072 blood plasma dataset uploaded to Pride as part of the work published in [24]. The dataset features 78 subjects evenly distributed between young (18-22 years) and old (55-65 years) groups. We randomly selected 50% of the subjects to prepare three prompt formats. In the binary classification format, LLMs were instructed to select whether a sample is old or young, based on the top-100 most expressed proteins in a profile with exact NPX values.

For the pairwise format, 380 prompts were prepared, in which an LLM was instructed to select the older sample, based on the top-100 most differentially abundant proteins. In the generative format, a model was presented with randomly sampled 25 proteins among the top-50 most highly expressed and instructed to generate the remaining 25. To increase the number of generative prompts, we split profiles into parts representing Olink’s categorization of sub-panels (Cardiometabolic, Inflammation, Neurology, and Oncology) and thus prepared four prompts from each sample. To account for a limited number of features measured by Olink platforms, we included all proteins a sub-panel could measure as part of the prompt. For all formats, control probes and probes below the limit of detection were excluded.

### 2.2. LLM benchmarking

We evaluated 15 LLMs across seven biomedical domains with tasks encompassing aging biology, survival analysis, and omics interpretation. The models tested included models developed by OpenAI (GPT-4.1, GPT-4.1-mini, GPT-5, GPT-5-mini, GPT-5.2, o3, o4-mini), Anthropic (Claude Sonnet 4.5), Google (Gemini 2.5 Pro, Gemini 2.5 Flash, Gemini 3 Pro Preview, Gemini 3 Flash Preview), xAI (Grok-3, Grok-4), and DeepSeek (DeepSeek-R1-0528).

Each task comprised multiple subtask formats designed to probe different aspects of model capability and assess the effect of different formats on model rankings. Classification tasks presented binary or multiple-choice questions, while pairwise comparison are a special case of a binary choice in which the labels represent not classes but entities, such as patient, individuals, or samples. Regression tasks required exact numerical predictions, while generative tasks required models to predict sets of genes characteristic of a given biological state (age and tissue). All prompts were constructed programmatically from source datasets and formatted with structured XML-like tags delineating the question, context, answer options, and ground truth. LLMs received task-specific system prompts instructing them to return structured JSON responses of an appropriate format. Models were queried through their respective APIs to Azure, Open Router, and xAI endpoints.

For unification purposes, the performance of LLMs in all binary, multiclass, and pairwise tasks is reported as unadjusted accuracy; in all generative tasks — as Jaccard index; in all regression tasks — as mean absolute error (MAE) in task-specific units.

## 3. Results

### 3.1. Benchmark dataset composition

We have aggregated data with various levels of annotation from seven disparate sources to compile a dataset that can both be used to test an LLM’s grasp of aging biology as well as fine-tune public models for aging-related research tasks.

The totality of the resulting prompt collection contains 30,193 prompts, which roughly corresponds to 50MM tokens, when tokenized with OpenAI’s tokenizer (see **Table 1**). In most cases, the number of prompts achieved has not exhausted all the information available in a source, meaning that an even more extensive collection may be compiled with these resources for supervised fine-tuning (SFT) and benchmarking purposes.

**Table 1.** Breakdown of the LongevityBench prompt collection by data source and biological domain. The LongevityBench prompts were prepared using seven independent data sources to display aging through the lenses of different biodata types. All tasks support multiple prompt formats and annotation depths to control semantic and contextual confounders.

| Task category | Primary source | Description | N prompts | N tokens |
| --- | --- | --- | --- | --- |
| General population survival | NHANES-IV | Given clinical blood tests and medical records of an individual, assess an individual's survival time. | 22,035 | 12,118 K |
| Masked gene aging | Open Genes | Given a description of a gene and its effects in model organism, establish its expression trajectory in a human tissue. | 648 | 349K |
| Multi-mutant lifespan | Synergy Age | Given the effect of one mutation on lifespan in a model organism, assess the effect of an additional mutation relative to wild type. | 192 | 76K |
| Cancer survival | TCGA | Given a patient's diagnosis and tumor mRNA expression data, assess progression free survival. | 1,634 | 863K |
| Methylation aging | GEO | Given a person's blood DNA methylation levels, predict their age. | 1,503 | 28,164 K |
| Transcriptomic aging | GTEX | Given a person's RNA expression levels, predict their age. | 3,603 | 7,976K |
| Proteomic aging | (Kirsher et al, 2025) | Given a person's Olink proteomics, predict their age. | 578 | 530K |
| TOTAL |  |  | 30,193 | 50MM |

### 3.2. Multi-task benchmarking

We evaluated 15 SotA LLMs across 17 LongevityBench tasks to establish a comprehensive ranking of their grasp of aging biology. The aggregate ranking, computed as the mean rank across all tasks, identified Google’s Gemini-3 Pro as the top-performing model with an average rank of 5.00, followed by Open.AI’s GPT-5 (5.06) and o3 (5.06). Despite high average scores achieved by some models (**Table 2**), there is major heterogeneity in model performance across data modalities. As such, no single model achieved top-3 placement across all tasks, and even the highest-ranked models display low results in certain domains (**Figure 2**).

**Table 2.**
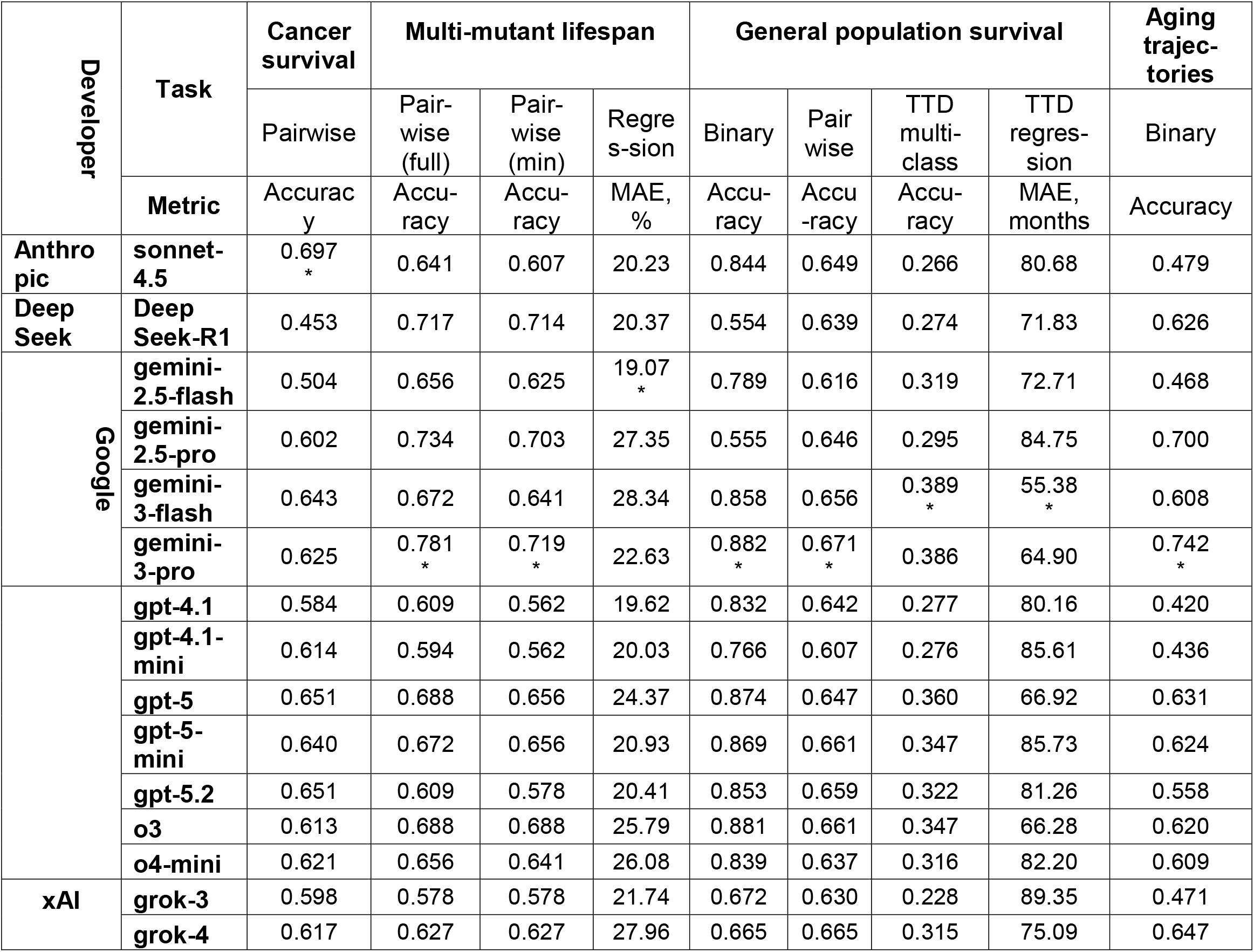
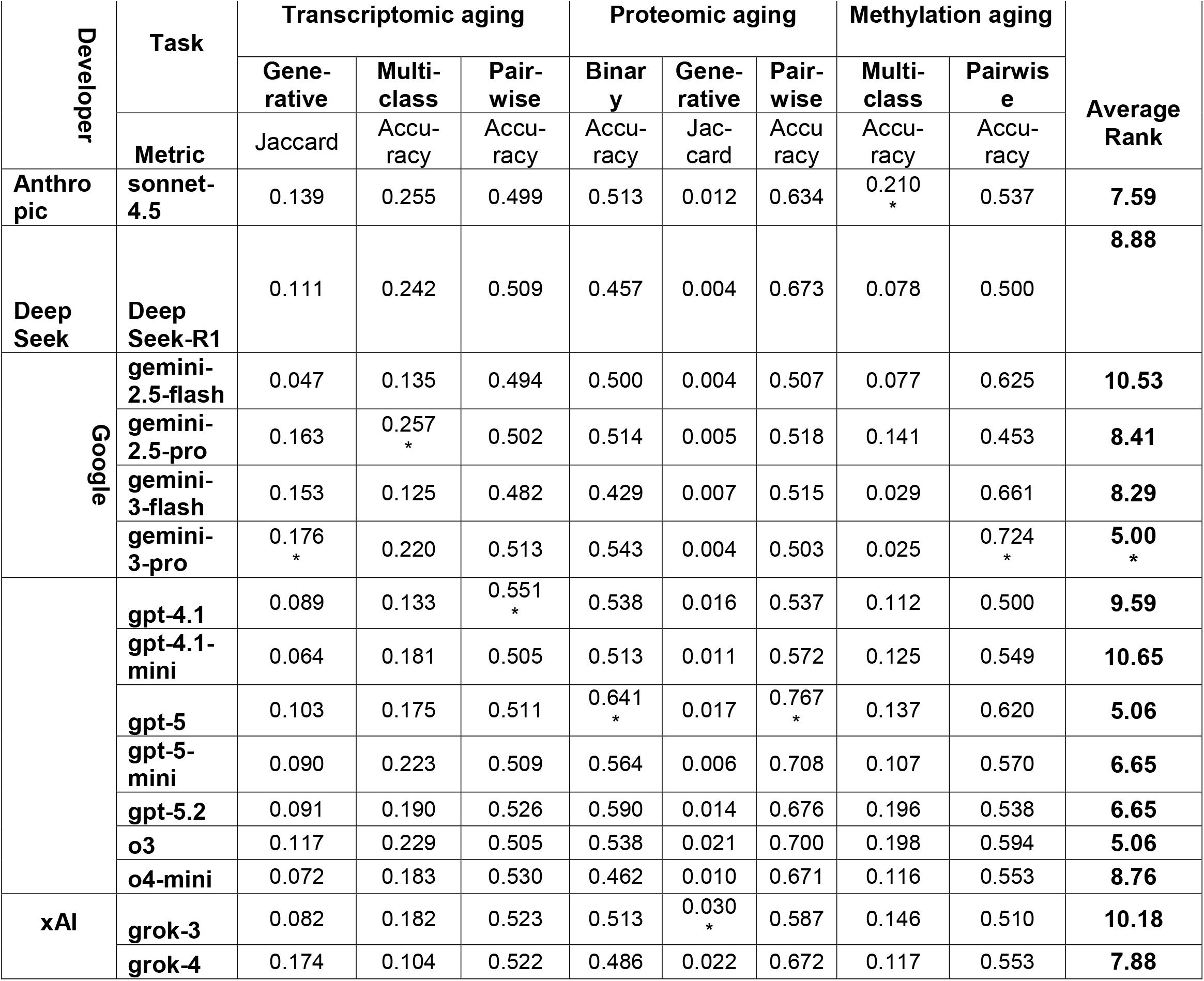
LongevityBench metrics across 17 tasks and 15 models.

**Figure 2.**
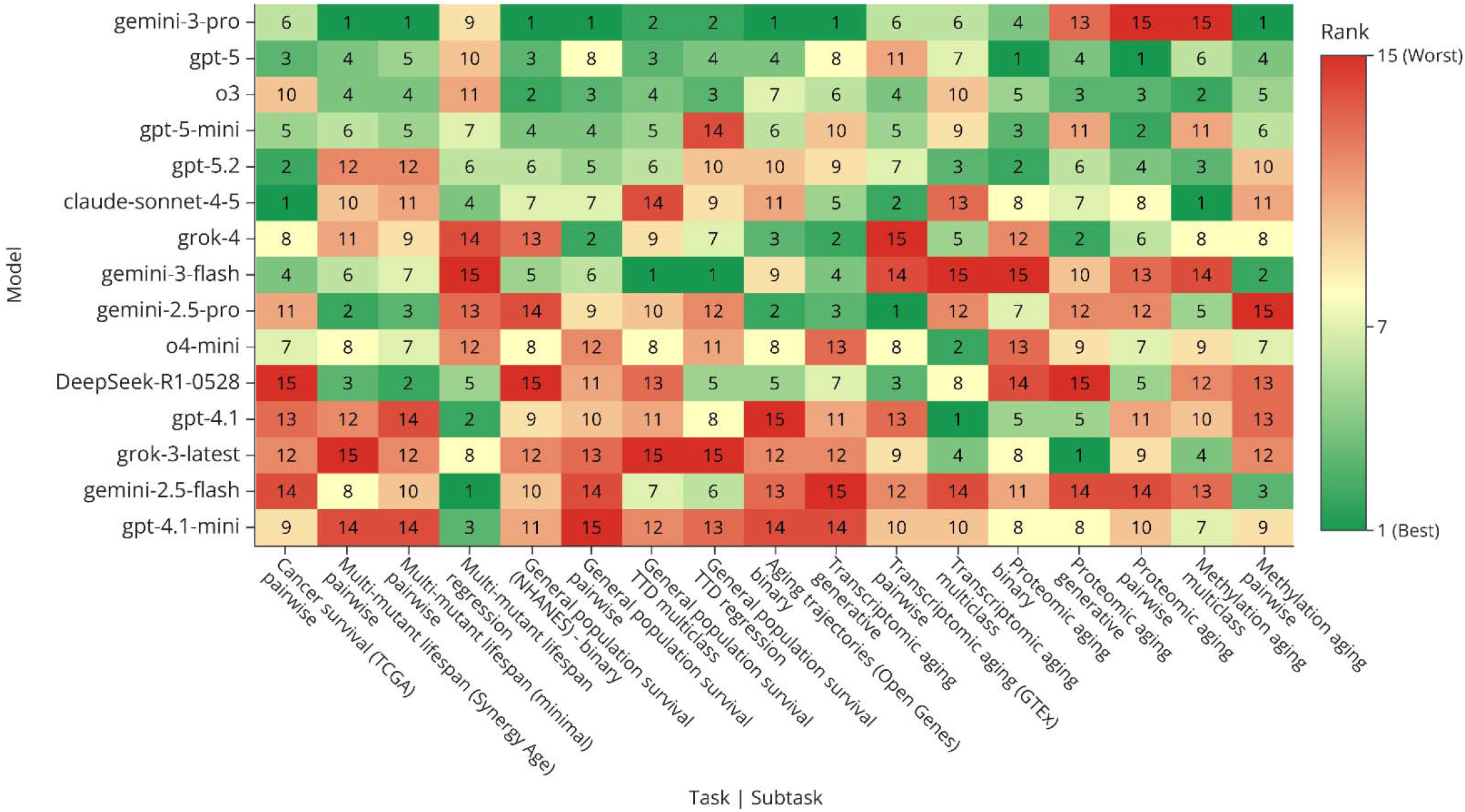
The ranks of 15 LLMs tested in 17 LongevityBench tasks. This collection of tasks represents common problems and settings encountered in aging research, such as survival analysis, omics-based age and lifespan prediction, determining the role of a gene in the aging process. The biodata domains covered by LongevityBench include: transcriptomics, epigenetics, proteomics, clinical blood tests, biometrics, and medical records. Rows ordered by an LLM’s average rank (see also **Table 2**).

The binary test results demonstrate that most models perform above the 0.5 random baseline, though margins vary considerably by task category. In the cancer survival task based on TCGA RNAseq data, Claude Sonnet 4.5 achieved the highest accuracy (0.697) when guessing which of the two patients had a longer PFS, with the close second being GPT-5.2 (0.651). For general population 10-year binary survival prediction based on NHANES clinical-level data, Gemini 3 Pro led with 0.882 accuracy, closely followed by o3 (0.881) and GPT-5 (0.874), only four models showed accuracy <0.7: xAI’s Grok-3 and 4, Gemini-2.5 Pro, and DeepSeek R1. Yet, in the pairwise setting using the same data (which individual lives longer?), all models showed much lower performance within the 0.607−0.671 range. The multi-mutant lifespan task, based on data from in murine and fly genetic experiments, had Gemini 3 Pro (0.781) and 2.5 Pro (0.734) taking the first two places, followed by Deepseek-R1 (0.717) (**Figure 3**).

**Figure 3.**
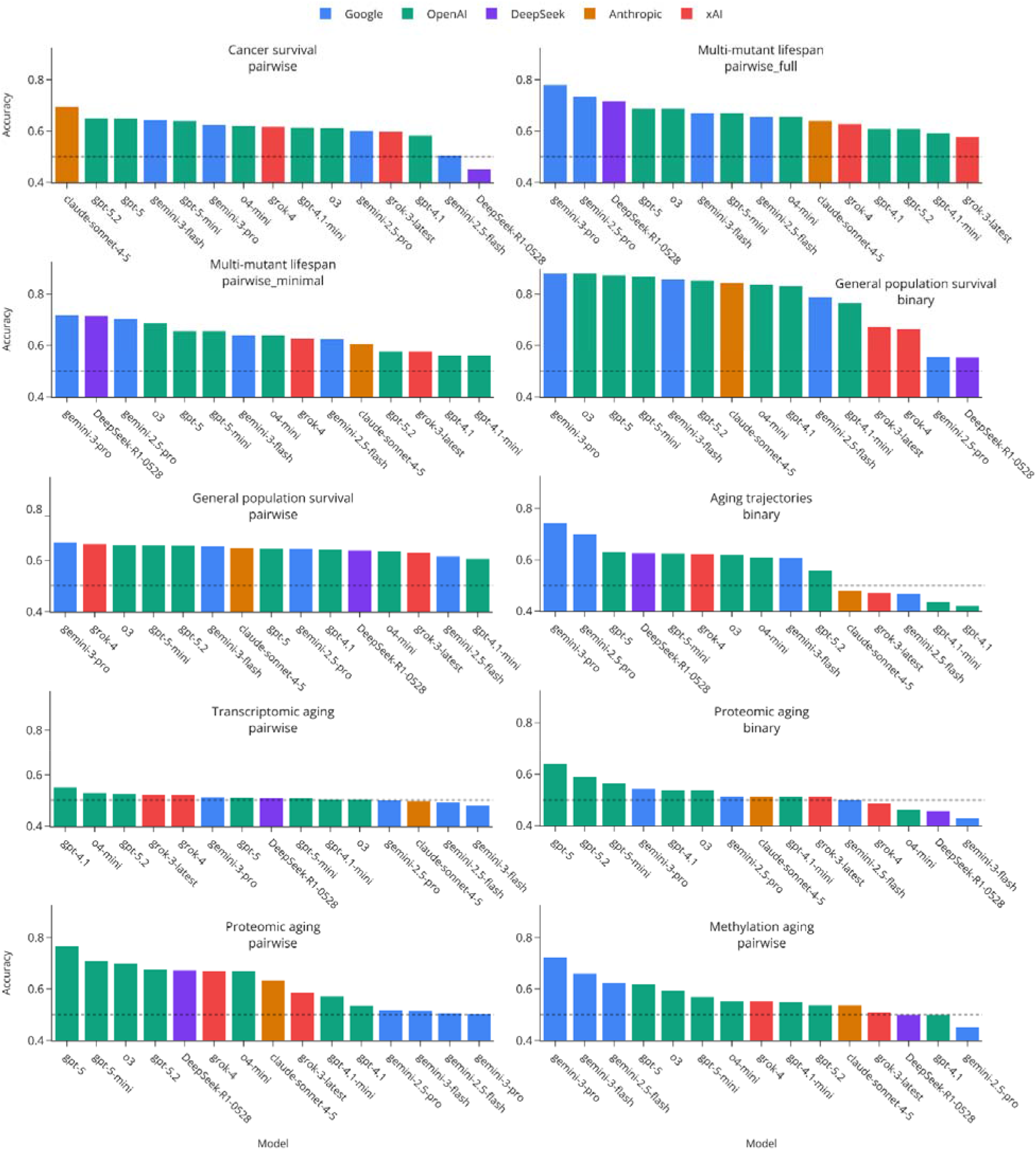
The performance of 15 tested LLMs in 10 tasks following a binary answer format. The presented tasks represent a variety problems encountered in aging research: survival analysis (“Cancer …” and “General population survival”), omics-based age prediction (“Transcriptomic …”, “Proteomic …”, “Methylation aging”), establishing the effect of a gene on longevity (“Multi-mutant lifespan”), and detecting an aging-related expression pattern (“Aging trajectories”). See **Methods** for the exact definition of each task.

The omics-level aging tasks presented greater challenges. In the transcriptomic pairwise age comparisons, we asked LLMs which of the two GTEx RNAseq profiles belonged to an older person, what we expected to be a relatively simple setting. However, all LLMs produced results indistinguishable from random baseline (0.482−0.551), suggesting that SotA LLMs struggle to extract aging signals from multi-tissue gene expression profiles. Notably, when the same data was presented as a multiclass age group prediction task (six decade bins covering the range of 20−79 years), LLM performance improved substantially: Gemini 2.5 Pro achieved 0.257 accuracy, well above the 0.167 random baseline. Furthermore, all developers except xAI had at least one model above the 0.20 accuracy threshold. This discrepancy indicates that aging-related transcriptomic patterns are encoded in these models, but the pairwise comparison format fails to elicit this knowledge. Proteomic aging prediction showed somewhat better discrimination in the pairwise format, with GPT-5 achieving 0.767 accuracy, though the binary classification (young vs old age groups) results for the same data type remained near random assignment level for most models with only GPT-5-mini, GPT-5.2, and GPT-5 showing accuracy above 0.55. In the epigenetic aging task, the pairwise format demonstrated that some SotA LLMs (Gemini and OpenAI families) can successfully detect aging signatures while others offered near-random predictions.

When all binary tasks are considered, models developed by OpenAI and Google dominate the leaderboard, with Gemini 3 Pro taking first place in 6 out of 10 binary tasks. Meanwhile, xAI models, Deepseek R1, and Anthropic’s Sonnet-4.5 generally show middling or low performance, with the notable exception of Sonnet’s top score in the cancer survival task (0.697 accuracy).

Since some tasks in LongevityBench are based on the same data sources, comparing LLMs based on the full array on 17 tasks may be unfair. To alleviate this bias toward models with a knack for overrepresented domains, we have selected one discriminant binary task per data source. Such tasks were selected to yield the greatest difference between minimal and maximal LLM scores. In this subset of tasks, Gemini-3 Pro shows outstanding performance compared to all others, but no single model dominates all domains. (**Figure 4**).

**Figure 4.**
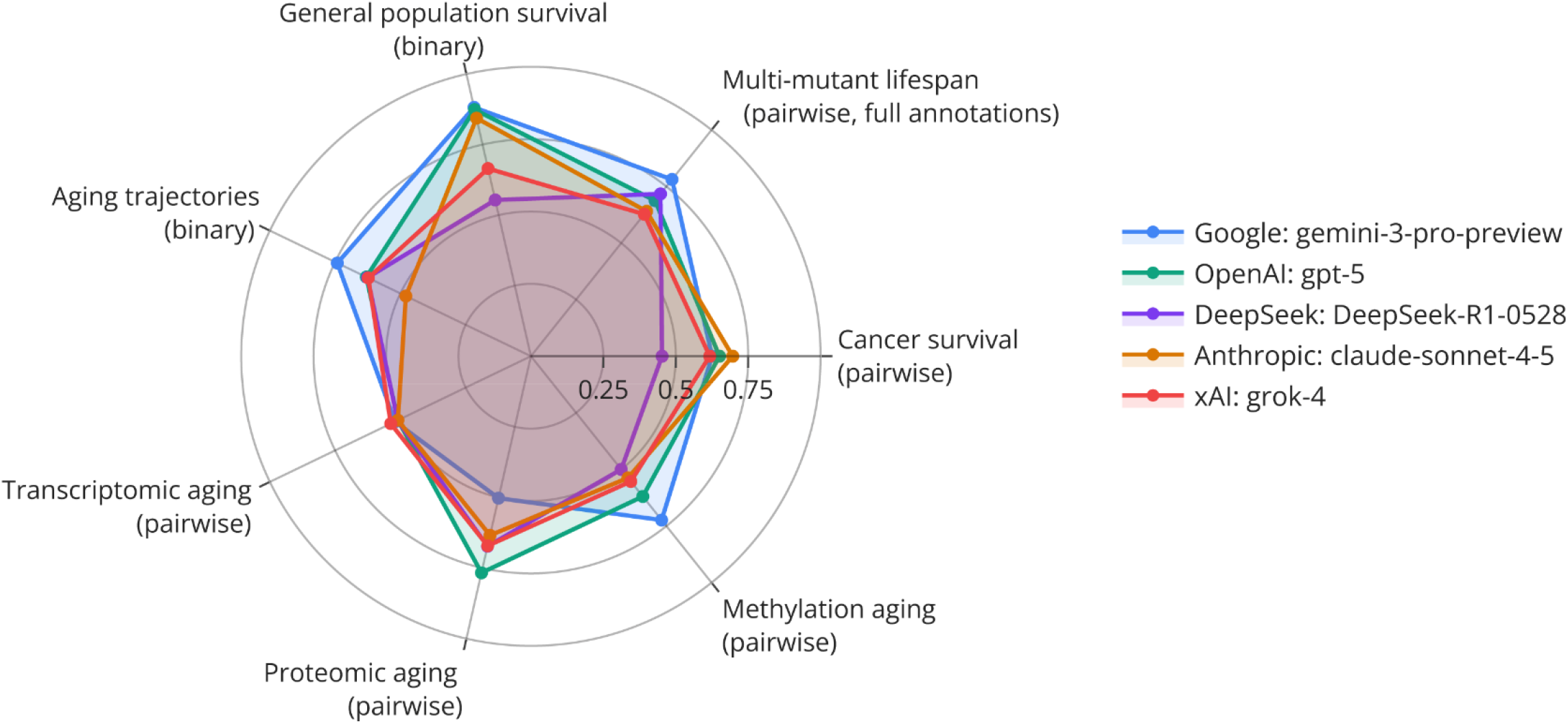
The comparison of best-performing LLMs from each developer across eight tasks selected for high inter-model variance. This tasks subset was defined to reduce the bias in model rankings stemming from the overrepresentation of certain data domains among the full set of 17 tasks.

### 3.3. Format-dependent variations

While using LongevityBench, we noticed that model rankings were not stable across question formats. We first examined this phenomenon using the multi-mutant lifespan task, where prompts were prepared in both minimal and extended annotation formats. The minimal format provided only the lifespan effects of individual mutations, while the extended format included GO annotations and experimental context from source publications. This comparison tests whether models can leverage additional biological context to improve phenotype-level predictions.

Several models demonstrated substantial improvement when provided with richer annotations: Gemini 3 Pro improved from 0.719 to 0.781, Gemini 2.5 Pro from 0.703 to 0.734. Similarly, GPT-4.1’s accuracy improved from 0.562 to 0.609, but the model with the best average rank, o3, remained stable at 0.688 in both conditions, suggesting that this LLM might have internalized to all the additional information we presented at training. Both tested xAI and DeepSeek-R1models showed no significant improvement as well. These patterns suggest that the capacity to extrapolate biological annotations into high-level phenotypes, such as longevity, varies across model families (**Figure 5**).

**Figure 5.**
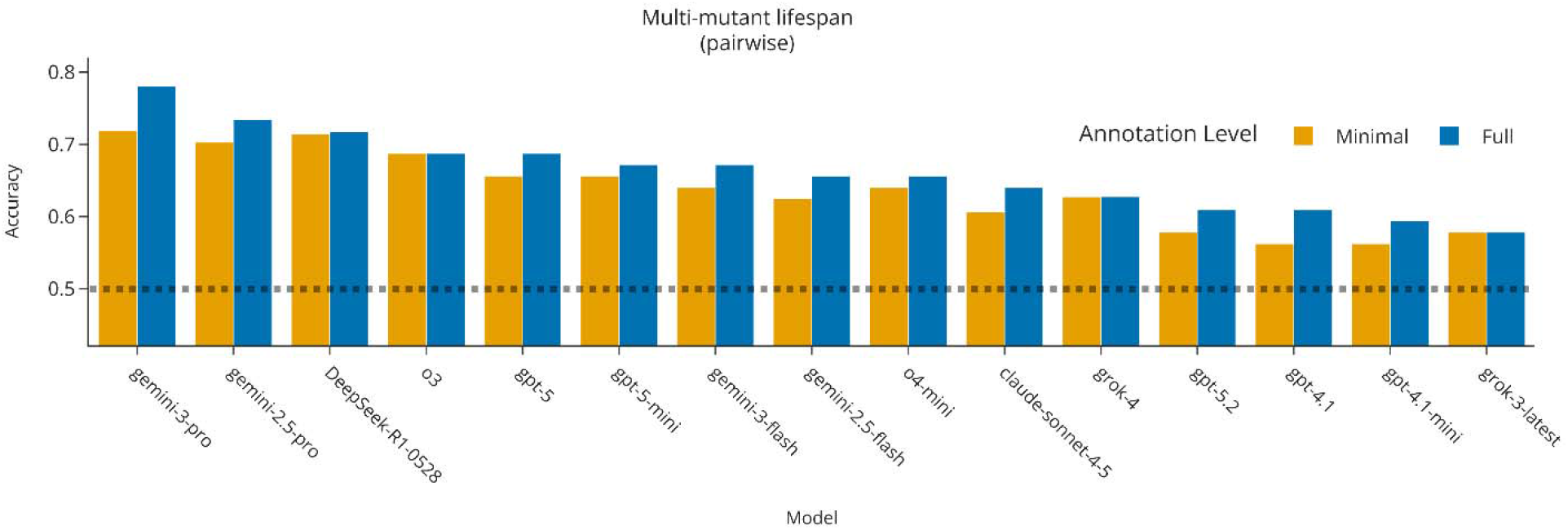
The change in LLMs’ performance in the “Multi-mutant lifespan” task caused by adding extra context to test prompts. Some models determine the longer-living genotype in *D*.*melanogaster* and *M*.*musculus* more accurately when presented with experiment protocols and descriptions of the affected genes.

The multiclass task formulations revealed additional format-dependent effects. In the NHANES survival prediction, the binned TTD format (four categories: 0−5, 5−10, 10−15, 15+ years) yielded different model rankings than the binary 10-year survival task using the same underlying data. Gemini 3 Flash achieved the highest multiclass accuracy (0.389), followed by Gemini 3 Pro (0.386), yet these same models ranked below random chance in the pairwise comparison format. Similarly, in methylation-based age prediction, Claude Sonnet 4.5 achieved the highest multiclass accuracy (0.210) despite middling performance in the pairwise setting (**Figure 6**).

**Figure 6.**
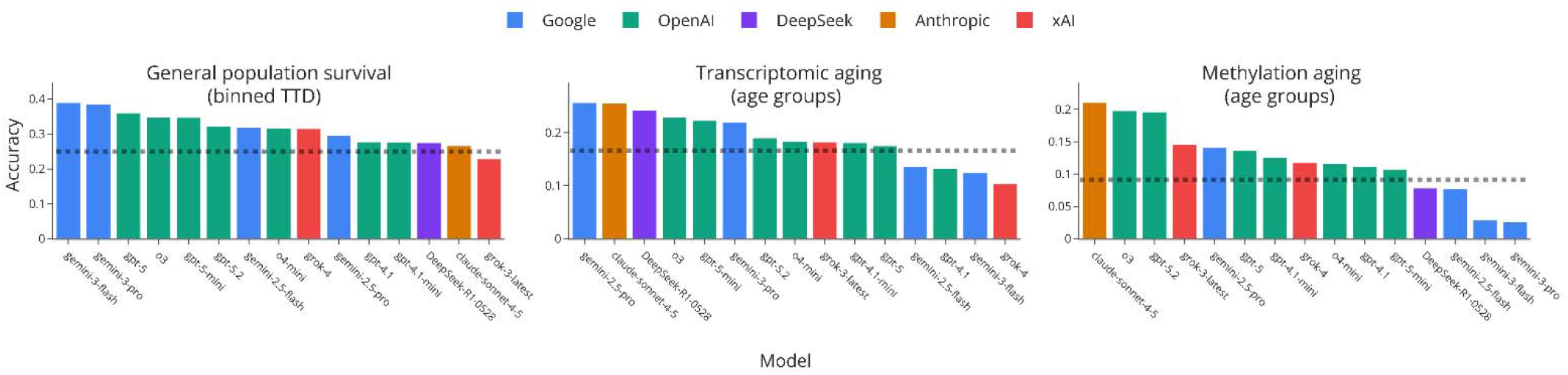
LLM performance in multiclass tasks from LongevityBench. Changing the format of the answer from pairwise choice (which sample is older or lived longer?) to multiclass age group prediction significantly affects LLM ranking (compare to **Figure 3**)

The regression tasks revealed systematic biases in model predictions. For multi-mutant lifespan effect prediction, models achieved MAE values ranging from 19.1% (Gemini 2.5 Flash) to 28.3% (Gemini 3 Flash), with most models showing reasonable regression parameters (**Figure 7**). However, the general population survival regression task exposed systematic underestimation: all models compressed their predictions toward the 50−100 month range, while true survival times extended beyond 250 months. This stands in stark contrast to the strong binary classification performance most models achieved on the same NHANES data, where accuracies exceeded 0.85 for top performers. The best regression model (Gemini 3 Flash, MAE 55.4 months) still exhibited this compression pattern, and weaker models (Grok 3 Latest, MAE 89.3 months) showed severe underestimation of long-term survival (**Figure 8**). The generative tasks, which required models to predict highly expressed genes given partial expression profiles and sample’s age and tissue metadata, revealed pronounced disparity between omics modalities. For transcriptomic aging, the top models achieved Jaccard similarities of 0.176 (Gemini 3 Pro) and 0.174 (Grok 4), indicating meaningful overlap with ground-truth gene sets. On the other hand, proteomic aging generation proved far more challenging despite a much narrower background set of measurable proteins. The best Jaccard similarity was only 0.030 (Grok 3), representing a six-fold reduction compared to transcriptomic performance (**Figure 9**).

**Figure 7.**
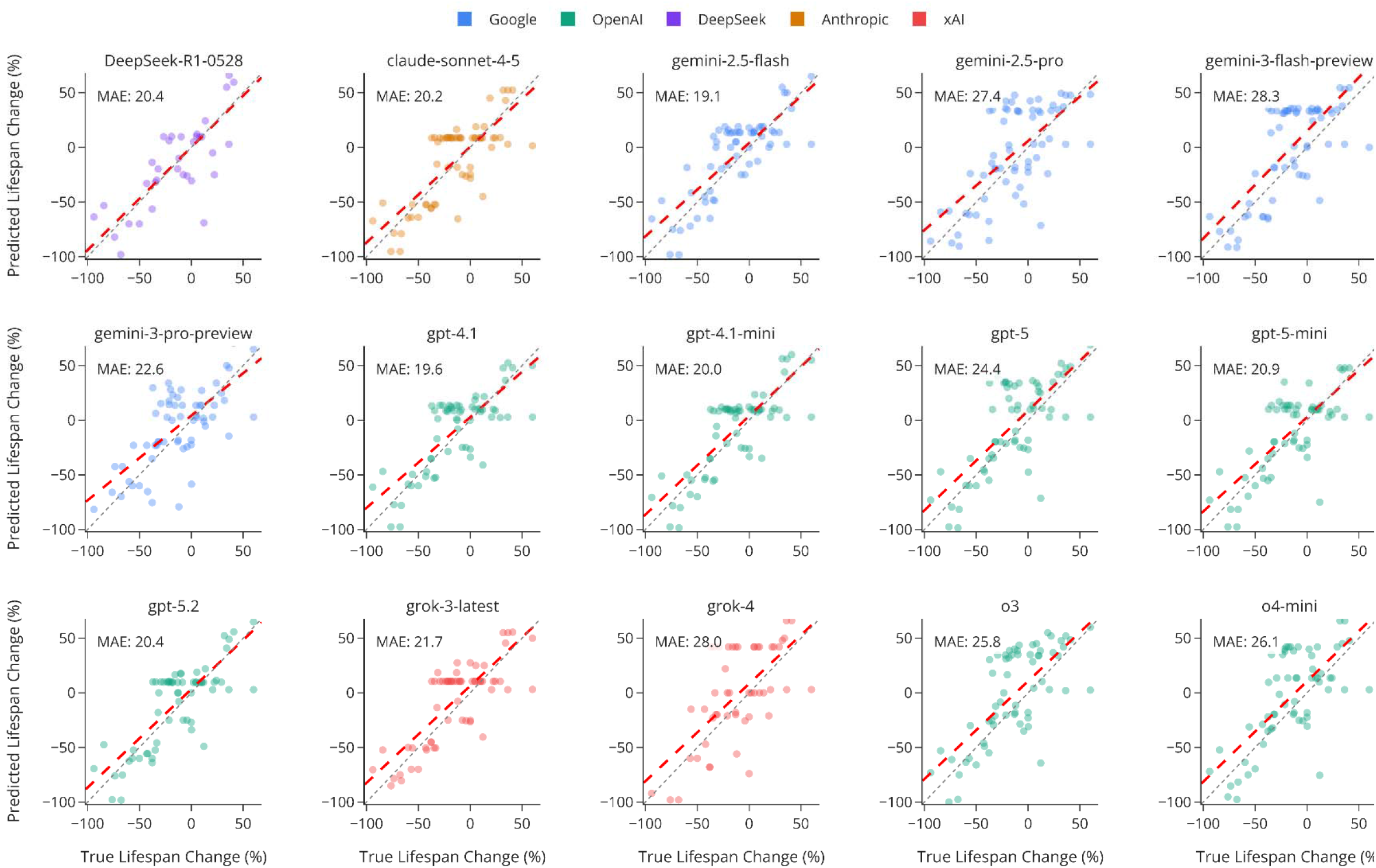
The performance of 15 LLMs in the “Multi-mutant lifespan” task which requires the models to provide the exact change in lifespan relative to wild type. The LLMs predict the combined effect of two mutations on model organisms’ lifespan with a 19-28% mean absolute error (MAE). Red line represents the ordinary least squares regression.

**Figure 8.**
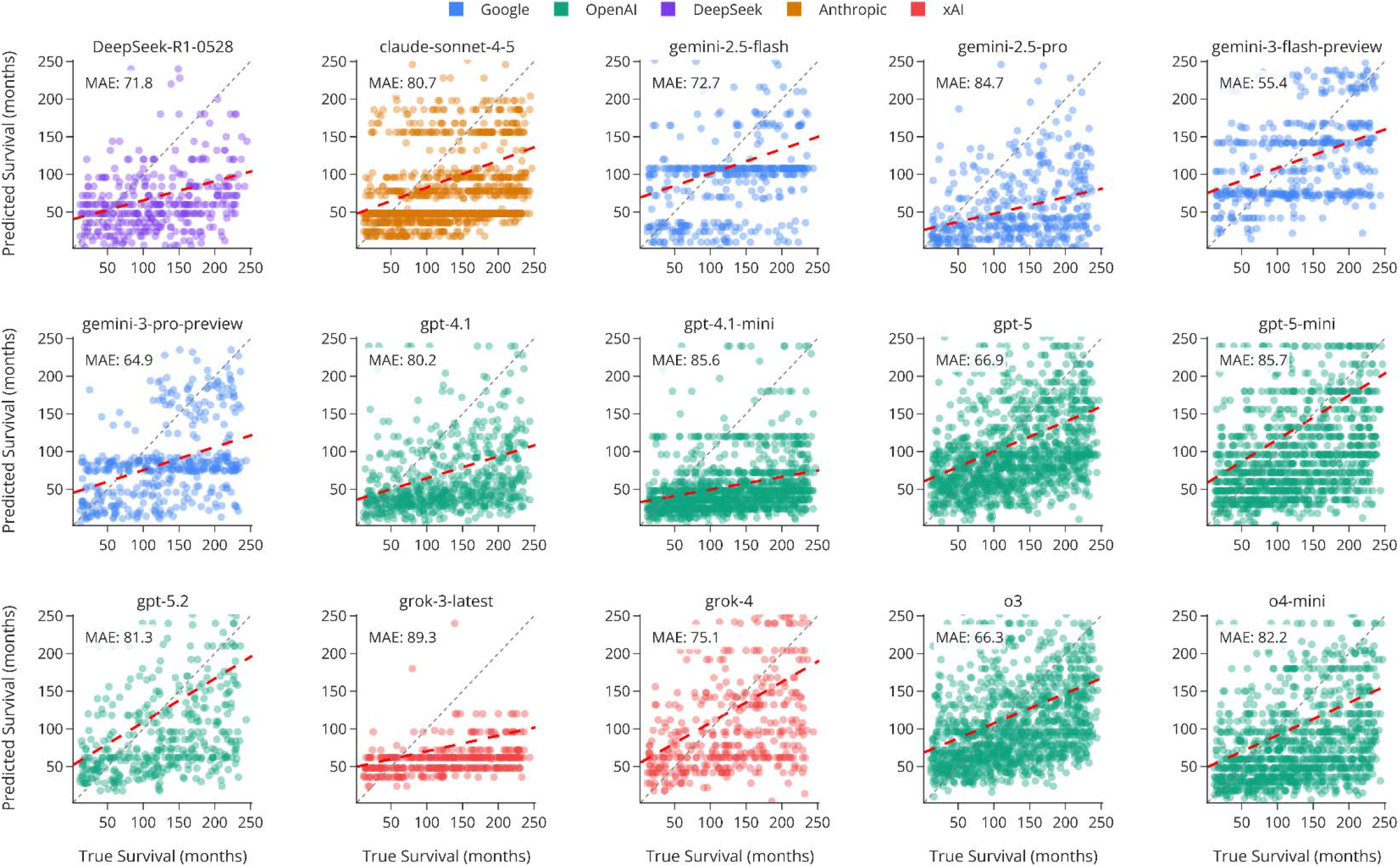
The performance of 15 LLMs in the “General population survival” task which requires the models to present the exact time-to-death (TTD) of an individual. Most LLMs tend to underestimate the TTD based on medical history and clinical blood tests, as indicated by the greater density of dots below the unity line (grey). Red line represents the ordinary least squares regression.

**Figure 9.**
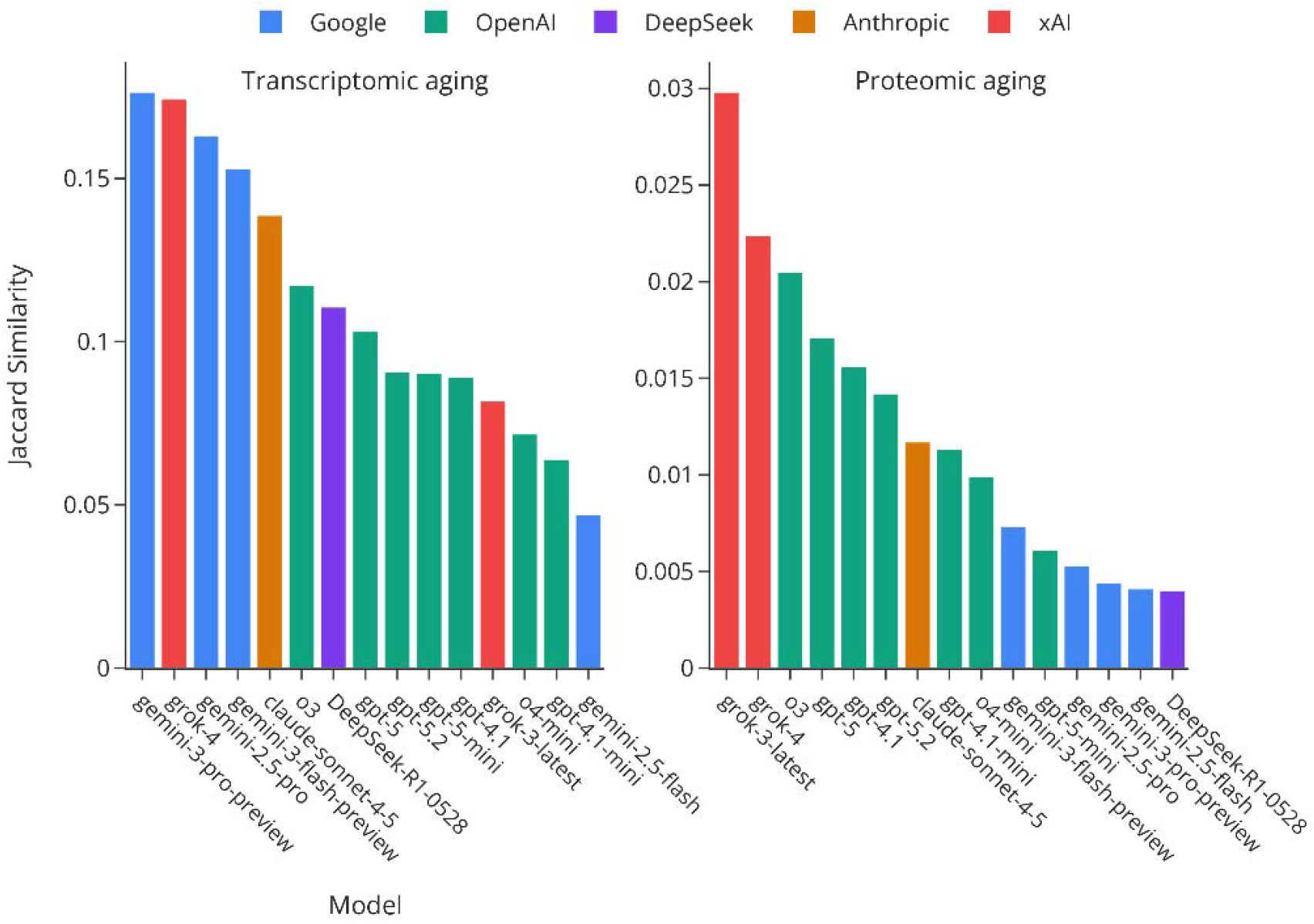
LLMs’ performance in generative omics tasks. Some LLMs can somewhat accurately complete lists of highly expressed genes in GTEx samples (Jaccard index > 0.15) but no LLM can do the same with Olink data (Jaccard index < 0.03) despite a much smaller background set of measurable genes.

## 4. Discussion

Our evaluation of 15 SotA LLMs reveals that no model demonstrates consistent strength across all modalities and question formats represented in LongevityBench. While individual models achieve strong performance on specific tasks, the substantial cross-task variance demonstrates that success in one domain does not guarantee competence in others (**Figures 2-4**). Researchers selecting LLMs for aging-related work must therefore consider specific task requirements rather than relying on general capability benchmarks or aggregate rankings. We note, however, that this assessment is incomplete and newer models, such as Claude Opus-4.5 or GPT-5.2 Pro, may perform differently. This benchmark thus requires ongoing assessment as models evolve.

When all tasks are considered, models developed by OpenAI and Google dominate the leaderboard, with Gemini 3 Pro achieving first place in 7 out of 17 tasks and the best average rank overall (5.0). GPT-5 and o3 tie for second position (average rank 5.06), with GPT-5 showing particular strength in proteomic tasks. Meanwhile, xAI models, DeepSeek-R1, and Anthropic’s Sonnet 4.5 generally show middling performance, with the notable exception of Sonnet’s top score in the cancer survival task (0.697 accuracy). Since some tasks in LongevityBench derive from the same data sources, comparing models across all 17 tasks may overweight certain domains. To address this, we selected one discriminative binary task per data source, choosing tasks that maximized inter-model variance. In this balanced subset, Gemini 3 Pro maintains its lead, but no single model dominates all domains (Figure 4).

LongevityBench evaluates three complementary capabilities: the interpretation of biological signal, robustness to variation in question format, and generalization across omics modalities. The benchmark does not directly measure mechanistic understanding of aging biology, which remains an open challenge that we aim to address in future versions. The deliberation of reasoning capabilities would require more thorough examination of the models’ thinking traces, human curation, and involvement of proprietary, unpublished datasets to eliminate conflating reasoning and retrieval. This benchmark version establishes baseline performance across task types and identifies systematic gaps that merit further investigation.

One such gap we report is the instability of LLM performance depending on the question format. While using the benchmark, we noticed that model rankings were not stable across different prompt collections representing the same data modality. We first examined this phenomenon using the multi-mutant lifespan task, where prompts were prepared with either minimal or extended annotations. The minimal format provided only the lifespan effects of individual mutations, while the extended format included GO annotations and experimental context from source publications.

This comparison tests whether models can leverage additional biological context to improve phenotype-level predictions. Several models demonstrated substantial improvement when provided with richer annotations: Gemini 3 Pro improved from 0.719 to 0.781, Gemini 2.5 Pro from 0.703 to 0.734. Similarly, GPT-4.1’s accuracy improved from 0.562 to 0.609. Notably, o3 remained stable at 0.688 in both prompt types, suggesting that this model may have internalized the additional biological context presented in the extended format during training. Both xAI and DeepSeek-R1 models showed no significant improvement as well. These patterns suggest that the capacity to integrate biological annotations into phenotype-level predictions varies across model families (**Figure 5**).

We then compared a different kind of format bias by changing the prediction setting between binary cutoff, multiclass, or pairwise comparison while maintaining the annotation richness level. In the NHANES survival prediction task, the binned TTD format (four categories: 0−5, 5−10, 10−15, 15+ years survival) yielded different model rankings than the pairwise task (which individual lived longer?) using the same underlying data. Gemini 3 Flash achieved the highest multiclass accuracy (0.389), yet this model ranked sixth in the pairwise comparison format. In a similar line, Grok 3 showed decent accuracy (0.630) in the pairwise NHANES task but performed below random chance (0.228) when the question type was changed to multiclass. We also noted that some question types are more illustrative of model disparity, as was demonstrated by a more uniform distribution of accuracies in the pairwise NHANES task, compared to the binary cutoff variant, which separated LLMs into different performance bands (**Figure 3**). The effect of the question semantics on performance is even more clearly seenin in methylation-based age prediction, where Claude Sonnet 4.5 achieved the highest multiclass accuracy (0.210) despite middling performance in the pairwise setting (**Figure 6**). Conversely, the three Gemini models acing the pairwise DNAm format moved to the bottom of the leaderboard in the binned age prediction task. We interpret this large variance in performance metrics and relative rankings across prompt variations as the absence of a coherent understanding of biology in the tested models. We argue that a model with an internal image of the aging process should rank at the top of the leaderboard despite semantic variations, annotation depth, or question type.

We also explored another type of LLMs’ bias with the regression question format. For multi-mutant lifespan effect prediction, models achieved MAE values ranging from 19.1% (Gemini 2.5 Flash) to 28.3% (Gemini 3 Flash), with most models showing reasonable regression parameters (**Figure 7**). However, the general population survival regression task exposed systematic underestimation: all models compressed their predictions toward the 50−100 month range, while true survival times extended beyond 250 months. This stands in stark contrast to the strong binary classification performance most models achieved on the same NHANES data (accuracy >0.85, **Figure 3**). The best regression model (Gemini 3 Flash, MAE 55.4 months) still exhibited this compression pattern, and weaker models (Grok 3 Latest, MAE 89.3 months) showed severe underestimation of long-term survival (**Figure 8**). We hypothesize this bias to stems from LLMs’ training corpora where clinical risk factors co-occur with mortality-related text. Models appear to treat any mention of serious disease as a substantial mortality risk, failing to appropriately weight information indicating successful treatment or long-term remission. We aim to further explore how sensitive LLMs are to red flags, such as family history of disease or past diagnoses, in this setting by inspecting model reasoning in future Longevity Bench experiments.

The generative tasks, which required models to predict highly expressed genes given partial expression profiles and sample metadata, revealed a major disparity between omics modalities. For transcriptomic aging, top models achieved Jaccard similarities of 0.176 (Gemini 3 Pro) and 0.174 (Grok 4), indicating meaningful overlap with ground-truth gene sets. Proteomic aging generation proved far more challenging despite a much narrower background set of measurable proteins: the best Jaccard similarity was only 0.030 (Grok 3 Latest) (**Figure 9**). This gap likely reflects the relative scarcity of proteomics data and reports in training corpora, partially attributed to the relative novelty of the Olink platform. Models may also incorrectly assume that protein abundance mirrors transcript levels, an assumption that fails for the post-transcriptionally regulated plasma proteins [25, 26]. These findings caution against applying LLMs to emerging data modalities without empirical validation.

The findings presented here offer domain-specific guidance for researchers considering LLM assistance. For clinical survival prediction from health records, most frontier models achieve reliable performance, though regression estimates should be interpreted cautiously given systematic compression toward shorter survival times. Omics-based age inference shows marked format dependence and researchers should test multiple question formulations before drawing conclusions based on LLM output. Proteomic tasks warrant particular caution since even top-performing models show dramatically reduced accuracy in this domain. Generative tasks requiring models to predict molecular profiles are not yet suitable for production use, though they may serve as hypothesis-generating tools when outputs are validated experimentally. In all cases, we recommend treating LLM outputs as preliminary rather than definitive, particularly for novel biological questions not well-represented in training data.

However, commercial LLMs undergo frequent updates, and the rankings reported here will inevitably change as providers release new model versions. We see LongevityBench’s primary value not in establishing a definitive model hierarchy, but in identifying systematic capability gaps that persist across model families. We will maintain updated leaderboards at insilicobench.insilico.com as new models become available, enabling the research community to track progress in the readiness of AI applications for aging research.

Furthermore, awareness of LLMs’ shortcomings allows us not only to practice healthy criticism toward AI-generated insights, but also guide model development to cover the identified gaps. The diversity of tasks spanning clinical records, multi-omics, genetic perturbations, and cross-species phenotypes creates training objectives that reward coherent biological representations over narrow pattern-matching. As a module within the MMAI Gym for Science framework (**Figure 1**), we envision LongevityBench as integral part of a broader ecosystem designed to advance AI capabilities in life sciences and aging research. The evaluation approaches demonstrated here thus establish a foundation for future work on integrating aging biology benchmarks into model development pipelines.

## 5. Limitations

This benchmark did not assess inference costs or latency, which remain relevant for high-throughput research pipelines. Some models require 30−50 seconds per response, and models with deep reasoning traces may consume substantially more tokens, creating barriers to large-scale deployment regardless of accuracy.

We note that model compliance with output format instructions varied, and failures to produce parseable responses were treated as missing data. While this approach avoids penalizing models for refusals, it obscures practical considerations: models that frequently deviate from specified formats complicate automation efforts. We aim to cover the metrics of execution time, costs, and format compliance rates in later iterations of LongevityBench.

The scope of tasks presented in this work does not cover all the settings encountered in aging research. We plan to continue growing the array of tests shown here to next include data from chemical screening and geroprotector experiments, cross-species translation, sequence-based, and other tasks.

## Supporting information

TIFF Figures

## 6. Resource availability

Current and historical leaderboard standings for all tasks and LLMs are available at insilicobench.insilico.com.

## 7. Abbreviations

AI: Artificial Intelligence
BMI: Body Mass Index
DNAm: DNA Methylation
GEO: Gene Expression Omnibus
GO: Gene Ontology
GTEx: Genotype-Tissue Expression
LLM: Large Language Model
MAE: Mean Absolute Error
MMAI: Multi-Modal AI
mRNA: Messenger Ribonucleic Acid
NES: Normalized Enrichment Score
NHANES: National Health and Nutrition Examination Survey
NPX: Normalized Protein eXpression
nTPM: Normalized Transcripts Per Million
PFS: Progression-Free Survival
RFT: Reinforcement Fine-Tuning
RNA: Ribonucleic Acid
RNAseq: RNA Sequencing
SFT: Supervised Fine-Tuning
SotA: State-of-the-Art
ssGSEA: Single-Sample Gene Set Enrichment Analysis
TCGA: The Cancer Genome Atlas
TTD: Time-to-Death
WT: Wild Type

## 8. Contributions

**AZ**: Conceptualization, Supervision, Resources, Writing – review & editing; **DS**: Formal analysis, Software, Validation; **VN**: Project Administration, Methodology; **SP**: Formal analysis, Validation; **DZ**: Formal analysis, Validation; **VA**: Project Administration, Resources, Supervision; **DU**: Investigation, Writing – review & editing; **AA**: Methodology, Project Administration, Supervision; **FG**: Conceptualization, Data Curation, Investigation, Methodology, Software, Visualization, Writing – original draft

## 9. Supplementary materials

### Prompt samples

#### General population survival

##### Binary

~~~
<question>
     Did this participant survive 10 years post-examination?
</question>
<options>
     A. Yes B. No
</options>
<annotation>
     <demographic>
          67-year-old male with college graduate or higher education of
          non-Hispanic white ethnicity rates health as excellent.
     </demographic>
     <biometric>
          BMI 25.1 kg/m2, height 177.7 cm, waist circumference 97.7 cm,
          body weight 79.3 kg, diastolic blood pressure 79.3 mm Hg,
          systolic blood pressure 130.7 mm Hg, pulse 51.3 bpm.
     </biometric>
     <blood>
          Albumin (g/dL) 4.30; Alkaline phosphatase (U/L) 94.00; Blood urea
          nitrogen (mg/dL) 11.00; …[TRUNCATED]… Vitamin B12 (pmol/L)
          261.25.
     </blood>
     <questionnaire>
          Rates own health as excellent. Family history of diabetes.
     </questionnaire>
</annotation>
<answer>
          A
</answer>
<follow_up>
     SEQN 11005: Alive at 18 years 5 months (censored)
</follow_up>
~~~

##### Pairwise

~~~
<question>
     Which participant had a longer survival period after examination?
</question>
<options>
     A. Participant A B. Participant B
</options>
<participant_a>
     <demographic>
          76-year-old male with less than 9th grade school education of
          non-Hispanic black ethnicity.
     </demographic>
     <biometric>
          BMI 29.1 kg/m2, height 160.5 cm, body weight 75.0 kg, diastolic
          blood pressure 60.0 mm Hg, systolic blood pressure 122.0 mm Hg,
          pulse 62.0 bpm.
     </biometric>
     <blood>
          Albumin (g/dL) 3.90; …[TRUNCATED]… Vitamin B12 (pmol/L)
580.07.
     </blood>
     <medication>
          Donepezil (taken for 4 years), Haloperidol (taken for 10 months).
     </medication>
     <questionnaire>
          Has medical history of arthritis; cancer.
     </questionnaire>
</participant_a>
<participant_b>
     …[TRUNCATED]…
</participant_b>
<answer>
     A
</answer>
<follow_up>
     Participant A (SEQN 18188): Deceased at 4 years 2 months, cause:
     Alzheimer’s disease. Participant B (SEQN 788): Deceased at 1 years 2
     months, cause: a heart condition
</follow_up>
~~~

##### TTD multiclass

~~~
<question>
     How long did this participant survive post-examination?
</question>
<options>
     A. 15+ years B. 10-15 years C. 5-10 years D. 0-5 years
</options>
<annotation>
     <demographic>
          80-year-old male with less than 9th grade of school education.
     </demographic>
     <biometric>
          BMI 24.2 kg/m2, height 165.3 cm, waist circumference 103.0 cm,
          body weight 66.2 kg, diastolic blood pressure 88.0 mm Hg,
          systolic blood pressure 170.0 mm Hg, pulse 82.0 bpm.
     </biometric>
     <blood>
          Albumin (g/dL) 4.30; …[TRUNCATED]… Vitamin B12 (pmol/L)
168.26.
     </blood>
     <medication>
          Beclomethasone (taken for 2 months), …[TRUNCATED]…
     </medication>
     <questionnaire>
          Rates own health as good. Has medical history of angina pectoris;
          stroke; curent asthma. Family history of heart attack.
     </questionnaire>
</annotation>
<answer>
     D
</answer>
<follow_up>
     SEQN 12418: Deceased at 1 years, cause: a heart condition
</follow_up>
~~~

#### Aging trajectories

##### Binary

~~~
<question>
     You are presented with a description of an unknown gene’s (“the gene”)
     function and experimental findings in model organisms and humans.
     In CD8+ T cells of humans between ages 23 and 65 years, the gene’s mRNA
     expression measured by microarray:
</question>
<options>
     A. increases with age B. decreases with age
</options>
<model_organisms>
     In the brain of rhesus monkeys between ages 5 and 31 years, the gene
     decreased expression by 43.0%, as established by measuring protein
     levels. …[TRUNCATED]…
</model_organisms>
<ppi>
     This gene interacts with CHML, CHML, KMO, and KMO in humans.
</ppi>
<go>
</go>
     This gene has molecular functions including 11-cis retinal binding,
     …[TRUNCATED]…
<protein_atlas>
     This gene mRNA (Tau=0.81, tissue-enhanced) is most expressed in
     cerebral cortex and liver (nTPM 6-11). …[TRUNCATED]…
</protein_atlas>
<answer>
     B
</answer>
<follow_up>
     Gene: OPN3. OPN3 showed decreased gene expression by 46.2% in CD8+ T
     cells between ages 23 and 65 years, measured by microarray. DOI:
     10.1111/j.1474-9726.2009.00534.x
</follow_up>
~~~

#### Multi-mutant lifespan

##### Regression

~~~
<question>
     What is the effect of the Cat(IF);Sod1(IF) genetic perturbation on the
     animal’s lifespan in % when compared to wild type?
</question>
<organism>
     Drosophila melanogaster
</organism>
<genes>
     Cat: Cat (Catalase) | Sod1: Sod1 (Superoxide dismutase 1)
</genes>
<go>
     Cat: Function: metal ion binding, oxidoreductase activity,
     …[TRUNCATED]… | Sod1: …[TRUNCATED]… Location: mitochondrial
     intermembrane space, peroxisome, cytoplasm
</go>
<experimental_context>
     Temperature: 25°C Diet: cornmeal-agar medium …[TRUNCATED]… Yeast
     FLP recombinase was used to allow induced overexpression of catalase
     and/or Cu/Zn-superoxide dismutase in adults. Expression of FLP
     recombinase was driven by the heat-inducible hsp70 promoter
     …[TRUNCATED]…
</experimental_context>
<known_effects>
     Cat(IF): −14.2% lifespan (39 days vs 45 days WT) | Sod1(IF): +27.8%
lifespan (60 days vs 47 days WT)
</known_effects>
<answer>
     +22.2%
</answer>
<follow_up>
     Cat(IF): 39 days (−14.2% vs 45 WT) [PMID:9858546] | Sod1(IF): 60 days
     (+27.8% vs 47 WT) [PMID:9858546] | Cat(IF);Sod1(IF): 57 days (+22.2% vs
     47 WT) [PMID:9858546] | Interaction: Opposite lifespan effects of
     single mutants
</follow_up>
<question>
~~~

##### Pairwise (full annotations)

~~~
<question>
     Based on the known effects of individual interventions, predict which
     of them two interventions will result in longer mean lifespan. Consider
     potential genetic interactions.
</question>
<options>
     A. Cat(IF);Sod1(IF) B. Cat(IF)
</options>
<organism>
     Drosophila melanogaster
</organism>
<genes>
     Cat: Cat (Catalase) | Sod1: Sod1 (Superoxide dismutase 1)
</genes>
<go>
</go>
     Cat: Function: metal ion binding, …[TRUNCATED]…| Sod1: Function:
     protein homodimerization activity, …[TRUNCATED]…
<experimental_context>
     Temperature: 25°C Diet: cornmeal-agar medium Fly stocks
     …[TRUNCATED]…
</experimental_context>
<known_effects>
     Cat(IF): −14.2% lifespan (39 days vs 45 days WT) | Sod1(IF): +27.8%
     lifespan (60 days vs 47 days WT)
</known_effects>
<answer>
     A
</answer>
<follow_up>
     …[TRUNCATED]…
</follow_up>
~~~

#### Cancer survival

##### Pairwise survival

~~~
<question>
     Which of the following two Melanoma cases has had a longer progression-
     free interval after the initial RNAseq screening:
</question>
<options>
     Patient-A: A 42-year-old female diagnosed with Stage IIIB ulcerated
     tumor (TT4b, NN1, MM0);
     Patient-B: A 77-year-old male diagnosed with Stage IIIC ulcerated tumor
     (TT4b, NN2b, MM0), Clark level IV
</options>
<gsea>
     Patient-A shows high activity in “Formation Of The Cornified Envelope”
     pathway (NES: 0.44, genes: KRT19, DSG2, KRT7, KRT18, KRT8 and 124 other
     genes), while Patient-B shows low activity (NES: −0.13, genes: PRSS8,
     DSG2, EVPL, PPL, DSP and 124 other genes). …[TRUNCATED]… Both
     patients show suppressed activity in the “Defective C1Galt1C1 Causes
     Tnps” pathway, with Patient-B demonstrating particularly strong
     enrichment (NES: −0.30, genes: MUC4, MUC15, MUC13, MUC20, MUC6 and 12
     other genes) compared to the other patient (NES: −0.15, genes: MUC13,
     MUC16, MUC6, C1GALT1C1, MUC17 and 12 other genes).
</gsea>
<answer>
     Patient-B
</answer>
<follow_up>
     Patient-A: TCGA-EB-A4IQ-01: Progressed after 11.1 months; Patient-B:
     (TCGA-GN-A26C-01: Progressed after 1.6 years)
</follow_up>
~~~

#### Methylation aging

##### Multiclass

~~~
<question>
     Based on the following DNA methylation profile from a blood sample,
     which age group does this individual most likely belong to?
</question>
<patient>
     A female from Denmark.
</patient>
<methylation_profile>
     Lowest methylated (75 sites): (chr12:121078658, + strand,
     CTTCAGCAGCGACGTCTTGGACTCGCTTTTGGCGATGTGCGAGCTCATCG): Located in CABP1
     (in first exon) (calcium binding protein 1 isoform 1, …[TRUNCATED]…
     enzyme inhibitor activity). Also known as: CALBRAIN; HCALB_BR Beta:
     0.01. | …[TRUNCATED]…
     ||
     Highest methylated (75 sites): (chr17:7833237, + strand, AATATTCAAGCTGCAGAGCCTTCAGCAGGCGGGGCTTATGCAGATGAGCG): Located in KCNAB3
     (within 1500bp of TSS) (potassium voltage-gated channel;
     …[TRUNCATED]… potassium ion transport). Also known as: AKR6A9;
     KCNA3B; KCNA3.1B; KV-BETA-3; MGC116886 Unclassified Cell type specific.
     Beta: 0.86. | …[TRUNCATED]…
</methylation_profile>
<options>
     A. 0-9 B. 60-69 C. 20-29 D. 100+ E. 80-89 F. 50-59 G. 90-99 H. 70-79 I.
     40-49 J. 10-19 K. 30-39
</options>
<answer>
     B
</answer>
<follow_up>
     True age: 62 years, age group 60-69 (GSM1506320, GSE61496).
</follow_up>
~~~

##### Pairwise

~~~
<question>
     Based on the following DNA methylation profiles from blood samples,
     which individual is older?
</question>
<options>
     A: A female from Estonia ; B: A female from United Kingdom, diagnosed
     with nephropathy and type 1 diabetes.
</options>
<methylation_data>
     The following CpG sites show the largest methylation differences
     between the two individuals: (chr1:247712591, + strand,
     CGCATTTGTAATGGTGAGGTACATTGTGCTTCTAGAAGGTTAAGCCTGAA): Located in
     C1orf150 (in gene body) (hypothetical protein LOC148823, synonyms:
     FLJ44728, RP11-978I15.8). Also known as: FLJ44728; RP11-978I15.8
     Patient-A: 0.74, Patient-B: 0.19. | …[TRUNCATED]…
</methylation_data>
<answer>
     B
</answer>
<follow_up>
     Patient-A: 25 years (GSM1425751, GSE59065). Patient-B: 39 years
     (GSM501660, GSE20067).
</follow_up>
~~~

#### Transcriptomic aging

##### Pairwise

~~~
<question>
     Which of the following two Skin samples comes from an older individual?
</question>
<options>
     A. Sample-A ; B. Sample-B
</options>
<sample_a>
     <demographic>
          Male
     </demographic>
     <expression>
          Top expressed genes: KRT1; KRT10; ND4; …[TRUNCATED]…
     </expression>
</sample_a>
<sample_b>
     <demographic>
          Male
     </demographic>
     <expression>
          Top expressed genes: FN1; COX1; COL1A2; …[TRUNCATED]…
     </expression>
</sample_b>
<gsea>
     Sample-A shows moderate activity in Formation Of The Cornified Envelope
     (NES: 0.18, genes: SPINK9, LCE4A, LCE3C, LCE3B, LCE3A and 124 other
     genes), while Sample-B shows very low activity (NES: −0.14, genes:
     LCE3A, KRT84, KRT40, LCE1D, LCE1E and 124 other genes).
     …[TRUNCATED]… Both patients show suppressed activity in Fatty
     Acids, with Sample-B demonstrating particularly strong enrichment (NES:
     −0.31, genes: CYP2A13, CYP2B6, CYP2F1, CYP4A11, CYP4F2 and 10 other
     genes) compared to the other patient (NES: −0.07, genes: CYP2F1,
     CYP2A13, CYP4F11, CYP2A7, CYP2B6 and 10 other genes).
</gsea>
<answer>
     A
</answer>
<follow_up>
     Sample-A (GTEX-148VJ-0626-SM-5LUAW): 70-79; Sample-B (GTEX-11WQK-0008-
     SM-5SI6T): 50-59
</follow_up>
~~~

##### Multiclass

~~~
<question>
     Based on this expression profile, predict the chronological age group of the individual.
</question>
<options>
     A. 70-79; B. 40-49; C. 50-59; D. 30-39; E. 20-29; F. 60-69
</options>
<context>
     Human Brain transcriptomics data (RNA-seq, log1p-transformed TPM). Sex:
male.
</context>
<expression>
     3388 genes: not detected |
     1076 genes (0.69-4.67): CSN1S1, DEFA8P, …[TRUNCATED]… |
     1079 genes (13.21-14.38): ANAPC5, CYC1 …[TRUNCATED]… |
     Top-200 expressed: ND4 (19.29), ND2 (19.20), …[TRUNCATED]…
</expression>
<answer>
     B
</answer>
<follow_up>
     Sample ID: GTEX-ZUA1-0011-R8b-SM-51MST; Actual age: 40-49
</follow_up>
~~~

##### Generative

~~~
<instructions>
     Given the sample metadata and 100 highly expressed genes, predict 100
     additional highly expressed genes from the same sample. Output HGNC gene symbols separated by semicolons (;). Output exactly 100
     genes.
</instructions>
<context>
     Tissue: Brain; Sex: female; Age group: 40-49
</context>
<input_genes>
     COX1; COX2; COX3; CYTB; SNAP25; …[TRUNCATED]…
</input_genes>
<answer>
     ND4; ATP6; ND2; ND1; CALM1; PKD1; …[TRUNCATED]…
</answer>
<follow_up>
     Sample ID: GTEX-12ZZX-0011-R11a-SM-5DUVJ; Age: 40-49
</follow_up>
~~~

#### Proteomic aging

##### Generative

~~~
<platform>
     Olink Explore 3072 proximity extension assay (PEA) plasma proteomics.
     Values are NPX (Normalized Protein eXpression, log2 scale).
     Cardiometabolic panel: Proteins pertinent to cardiovascular and
     metabolic disease biology, providing broad coverage of relevant
     pathways and processes. Includes both well-established biomarkers and
     more exploratory potential markers. | 729 measured proteins:
     AAMDC;ABCA2; …[TRUNCATED]…
</platform>
<question>
     This plasma proteomics sample is from a 55–65 year old M. Given 25 of
     the top-50 most abundant proteins (NPX), predict the remaining 25 (in
     any order).
</question>
<olink>
     GLO1 (5.70); KIF22 (4.63); RANBP1 (4.17); EHD3 (3.68);
…[TRUNCATED]…
</olink>
<answer>
     CSRP3;MYH9;TNNI3;…[TRUNCATED]…
<follow_up>
     Subject: 5530211349; Age: 55–65; Sex: M
</follow_up>
~~~

##### Pairwise

~~~
<question>
     Based on their plasma proteome profiles, which individual is older?
</question>
<options>
     A. Patient-A B. Patient-B
</options>
<platform>
     Olink Explore 3072 proximity extension assay (PEA) plasma proteomics.
     Values are NPX (Normalized Protein eXpression, log2 scale). 2916
     proteins measured across Cardiometabolic, Inflammation, Oncology, and
     Neurology panels.
</platform>
<olink>
     Top-100 differentially abundant proteins (NPX): CASP1 (A: −8.89, B: −
     0.74); FOLR3 (A: 3.61, B: −4.33); …[TRUNCATED]…
</olink>
<answer>
     B
</answer>
<follow_up>
     Patient-A: 5530211325, 18–22, F; Patient-B: 5530211357, 55–65, M
</follow_up>
~~~

##### Binary

~~~
<platform>
     Olink Explore 3072 proximity extension assay (PEA) plasma proteomics.
     Values are NPX (Normalized Protein eXpression, log2 scale). 2916
     proteins measured across Cardiometabolic, Inflammation, Oncology, and
     Neurology panels.
</platform>
<question>
     Based on the plasma proteome profile of this male individual, what is
     their age group?
</question>
<olink>
     Top-100 most abundant proteins (NPX): GLO1 (5.67); BANK1 (5.07);
     …[TRUNCATED]…
</olink>
<options>
     A. 18–22 B. 55–65
</options>
<answer>
     B
</answer>
<follow_up>
     Subject: 5530211502; Age: 55–65; Sex: M
</follow_up>
~~~

