## Supplementary figures and images for "LongevityBench: Are SotA LLMs ready for aging research?"

### Figure 1.jpg

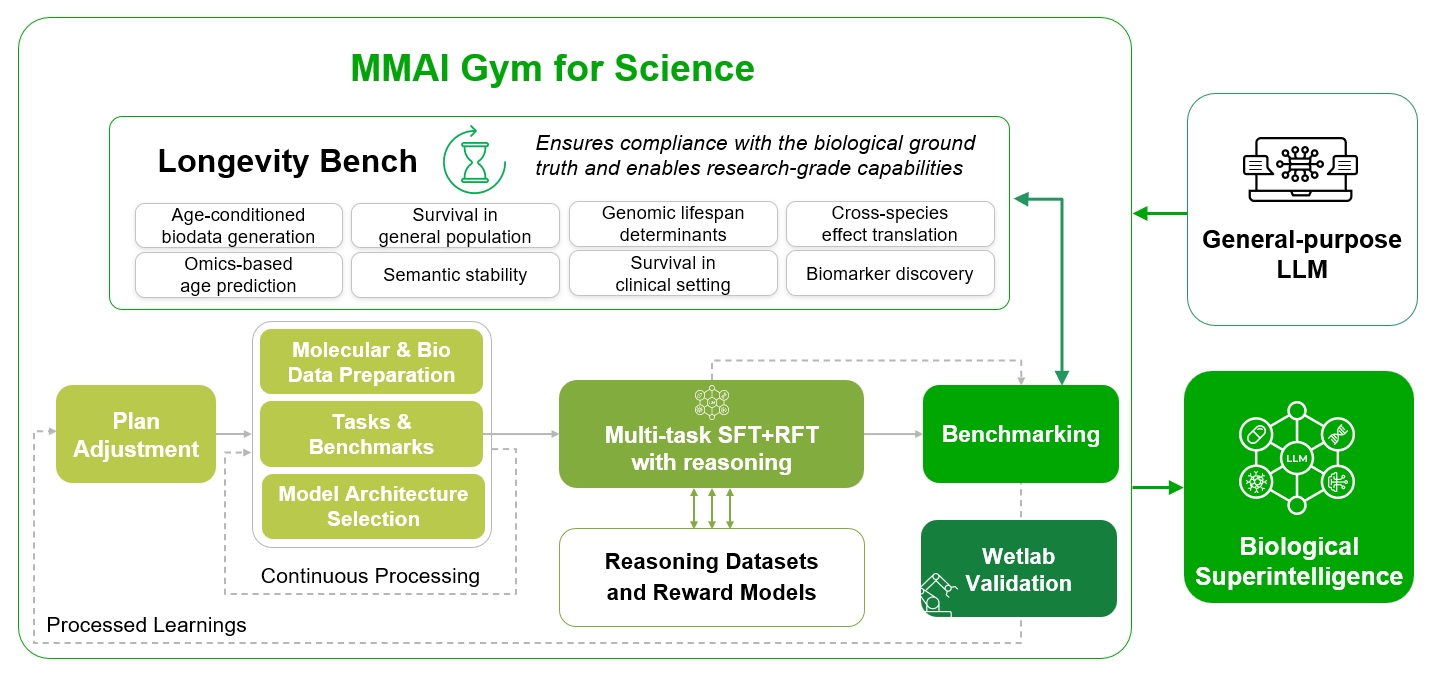

### Figure 2.tif

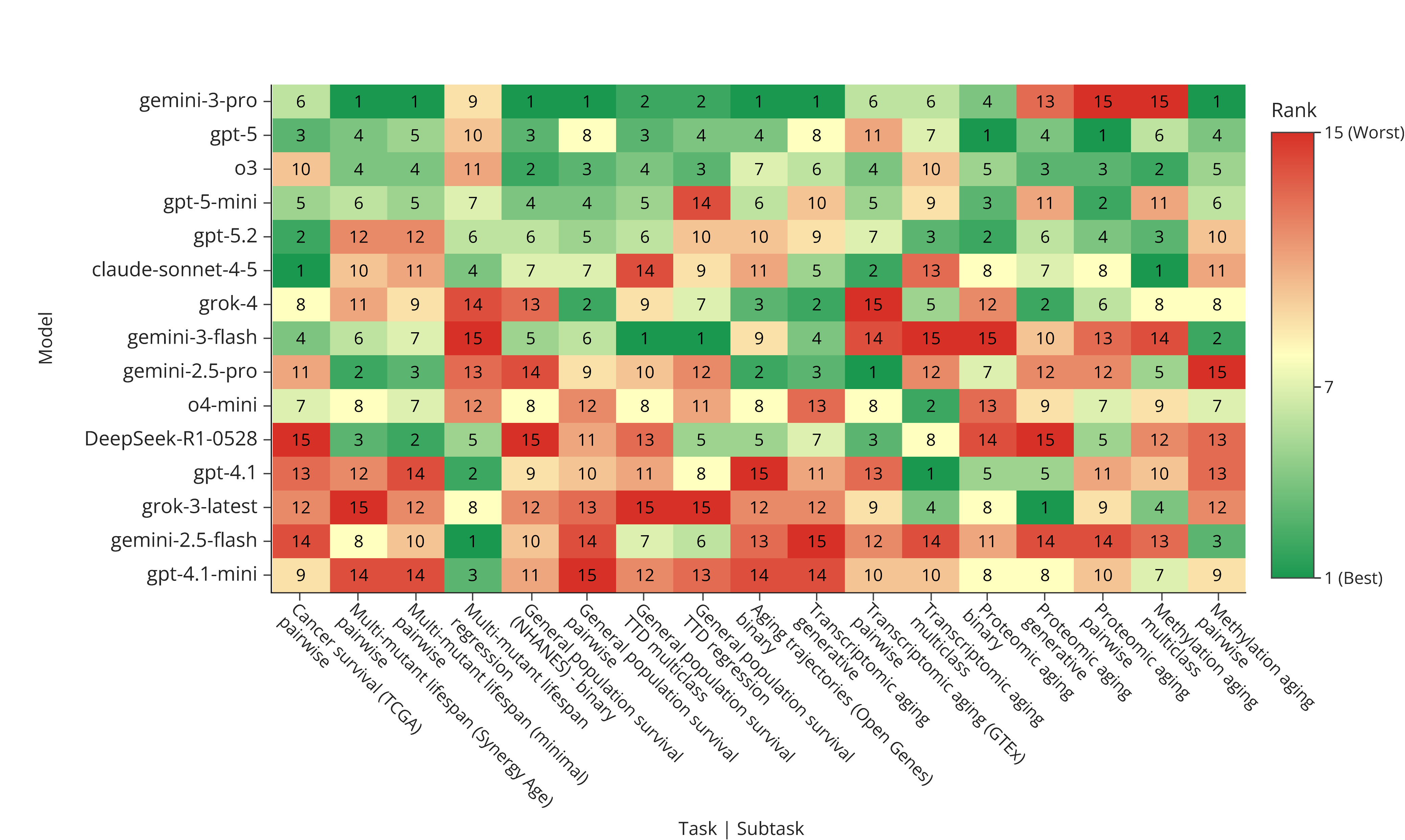

### Figure 4.tif

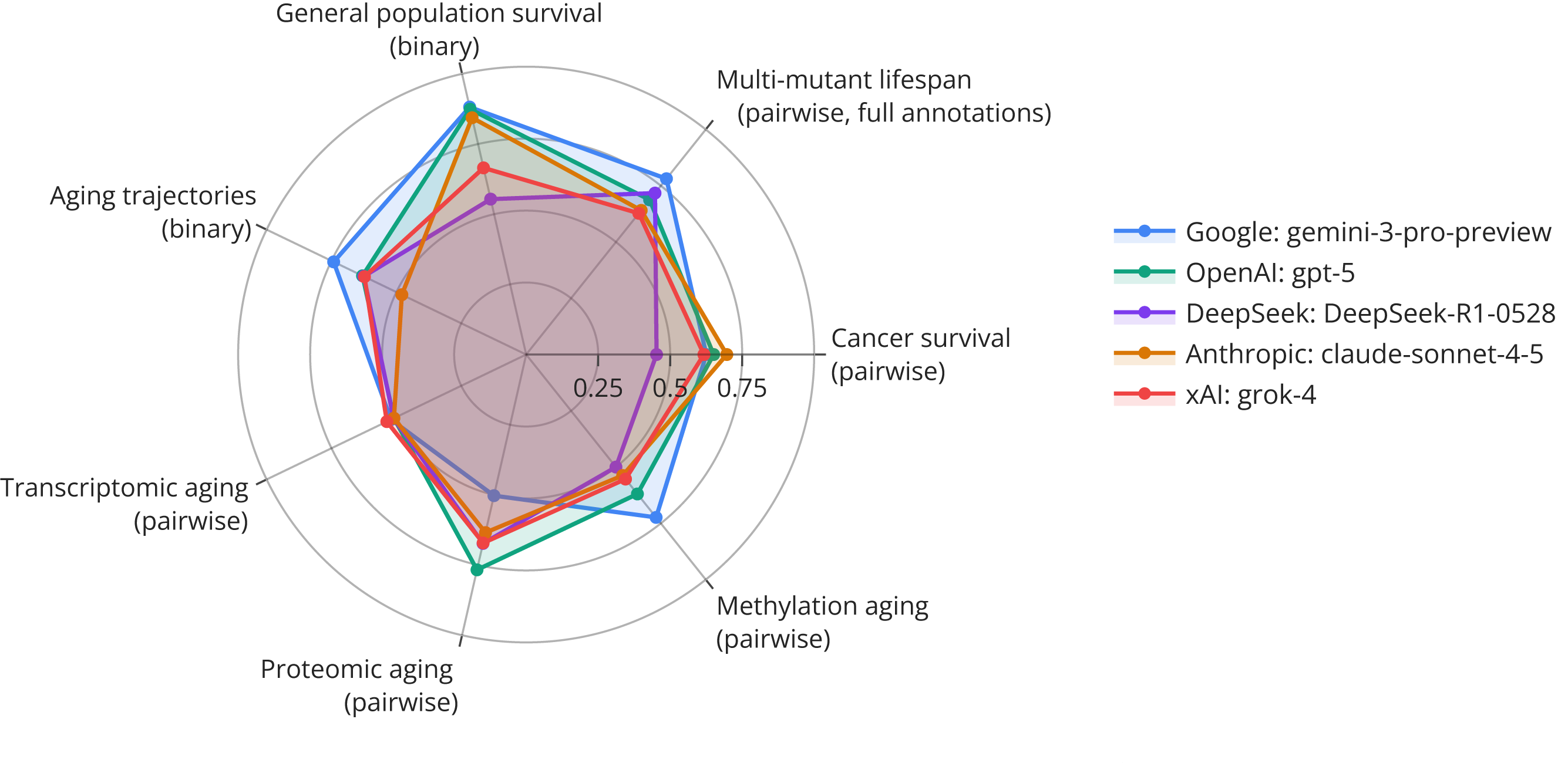

### Figure 5.tif

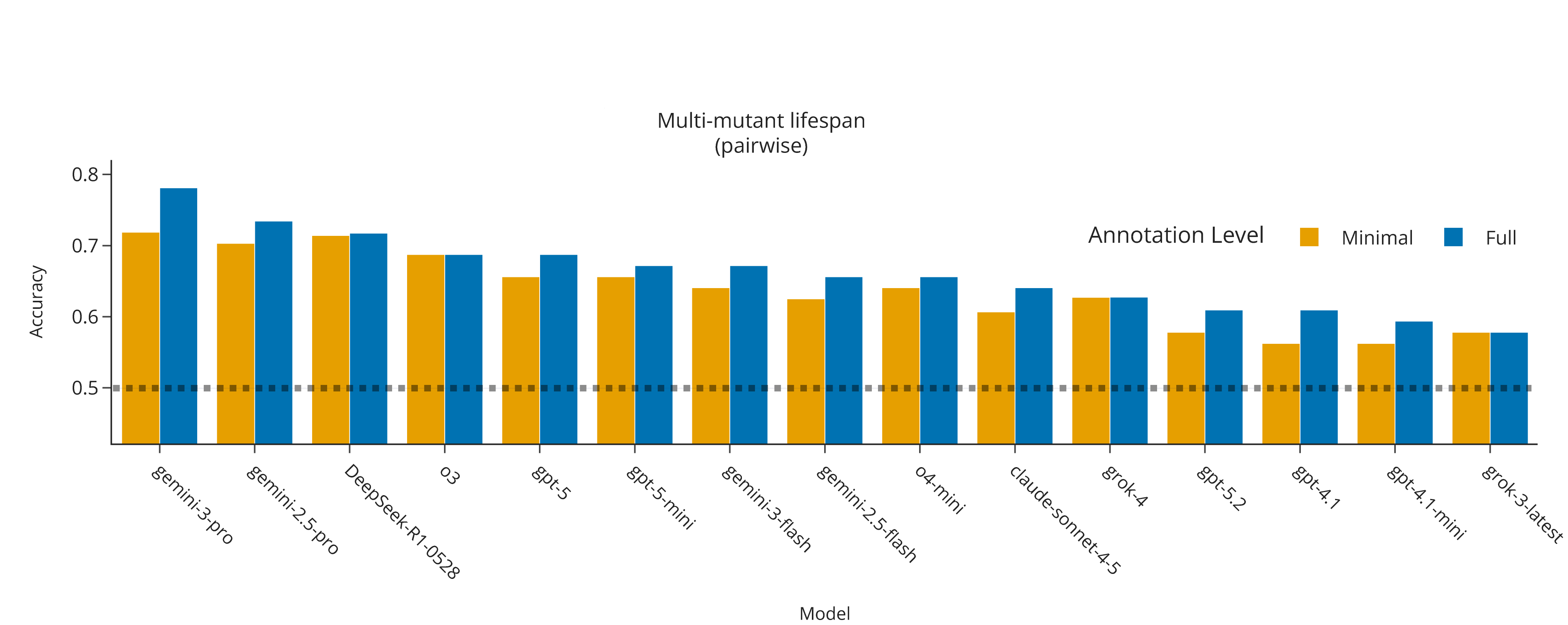

### Figure 6.tif

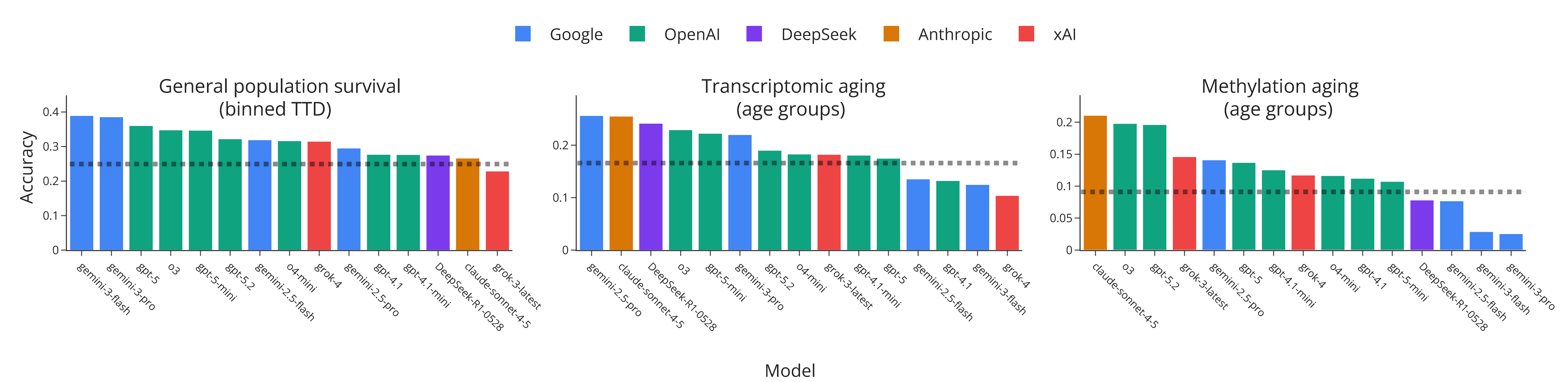

### Figure 7.tif

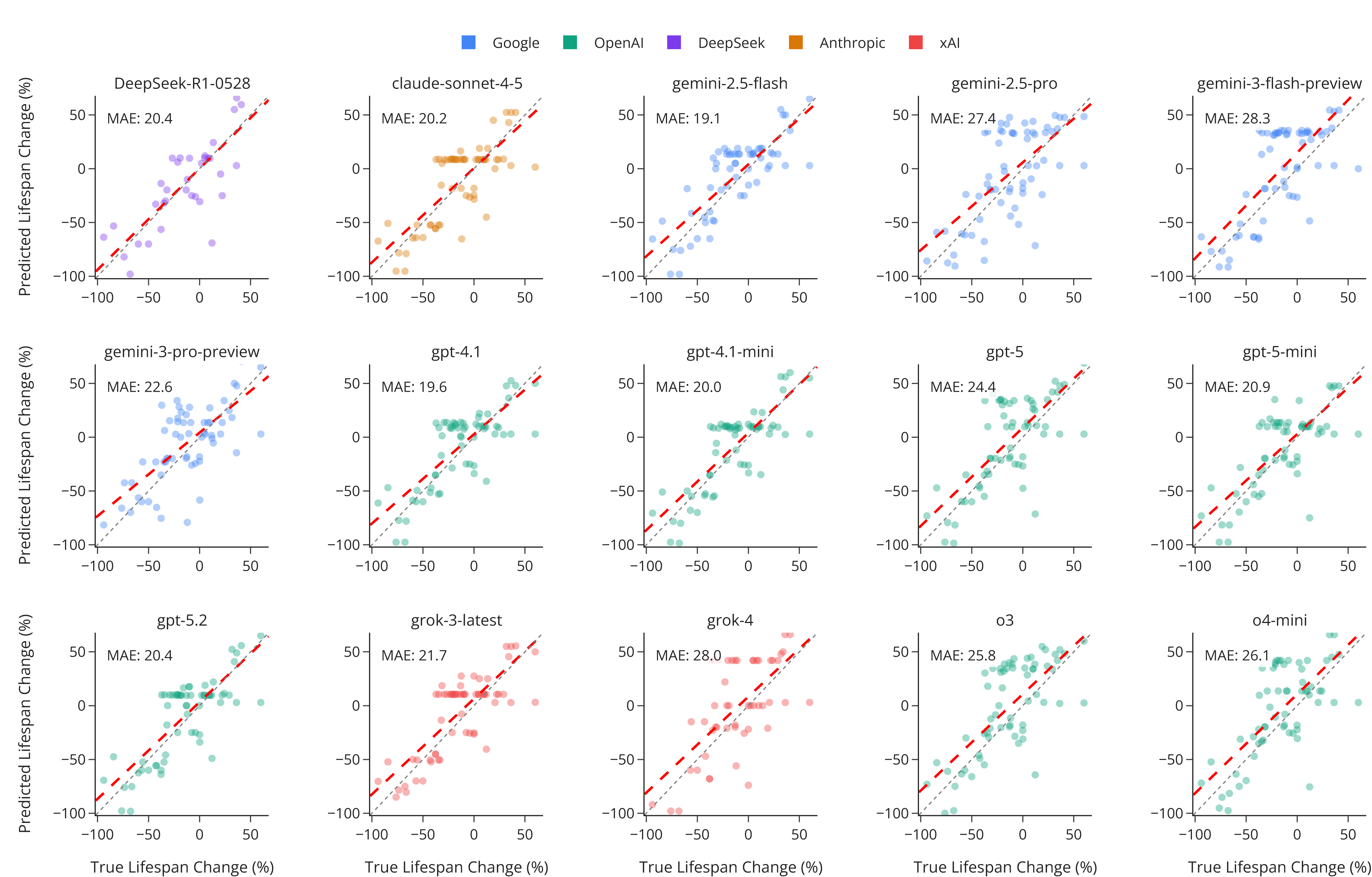

### Figure 8.tif

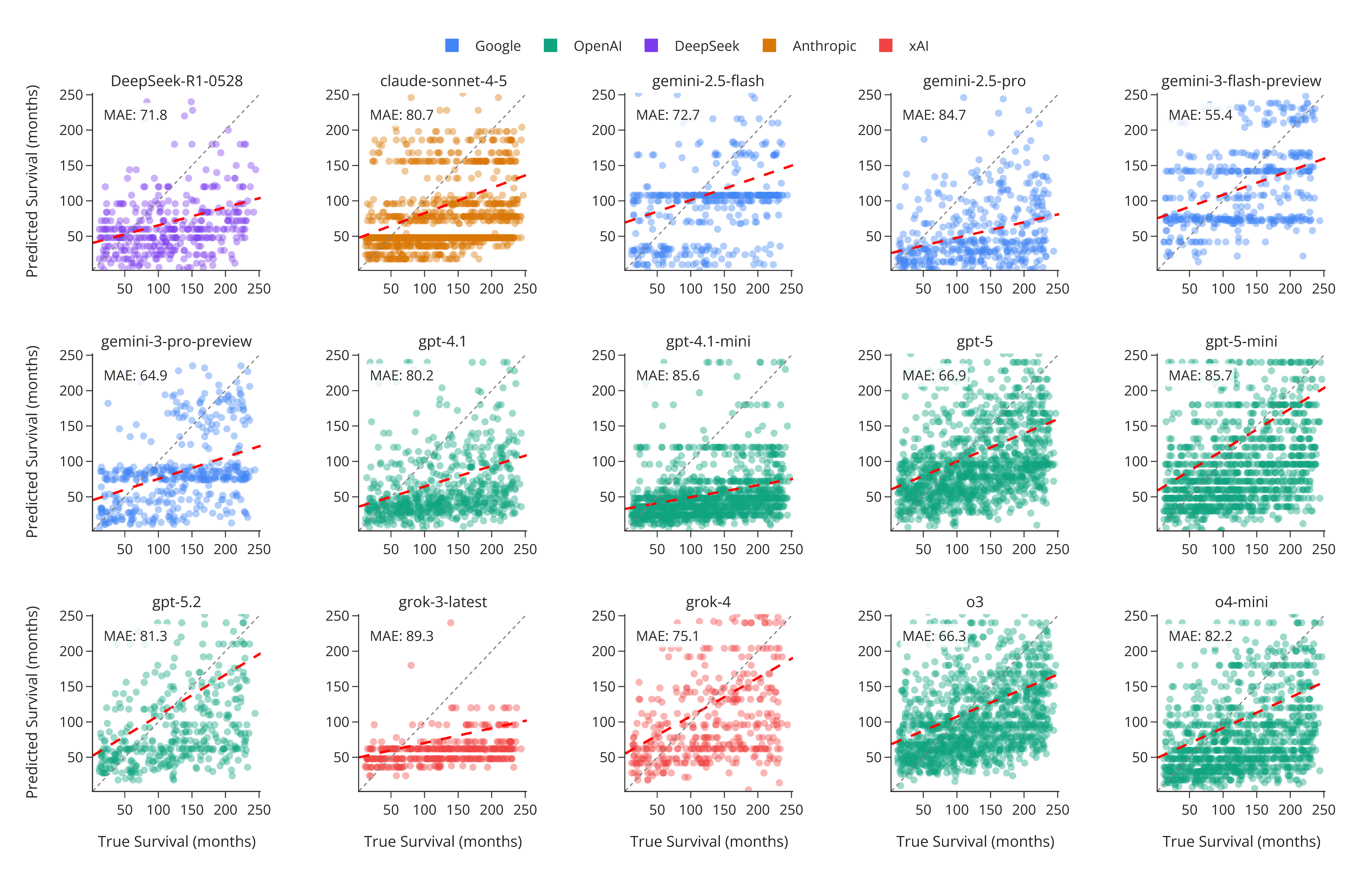

### Figure 9.tif

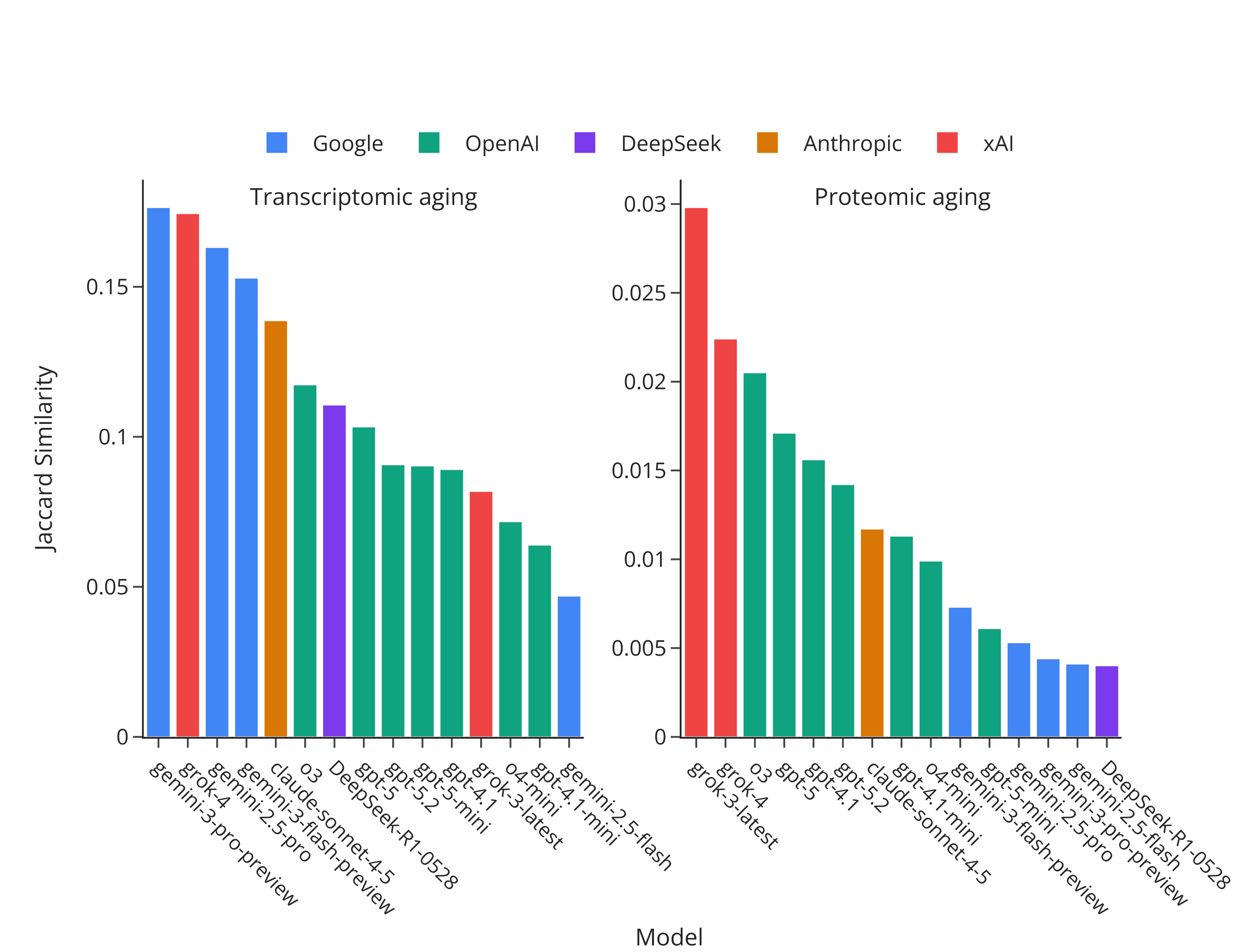
